# Exploring functional conservation *in silico*: a new machine learning approach to RNA-editing

**DOI:** 10.1101/2023.11.21.568001

**Authors:** Michał Zawisza-Álvarez, Jesús Peñuela-Melero, Esteban Vegas, Ferran Reverter, Jordi Garcia-Fernàndez, Carlos Herrera-Úbeda

**Author notes:** Contributed equally (Jesús Peñuela-Melero, Esteban Vegas, Ferran Reverter). **Materials & Correspondence.** Jordi Garcia-Fernàndez & Carlos Herrera-Úbeda.

## Abstract

Around 50 years from now, molecular biology opened the path to understand changes in forms, adaptations, complexity, or the basis of human diseases, through myriads of reports on gene birth, gene duplication, gene expression regulation, and splicing regulation, among other relevant mechanisms behind gene function. Here, with the advent of big data and artificial intelligence (AI), we focus on an elusive and intriguing mechanism of gene function regulation, RNA editing, in which a single nucleotide from an RNA molecule is changed with a remarkable impact in the increase of the complexity of transcriptome and proteome. We present a new generation approach to assess the functional conservation of the RNA-editing targeting mechanism using two AI learning algorithms, random forest (RF) and bidirectional long short-term memory (biLSTM) neural networks with attention layer. These algorithms combined with RNA-editing data coming from databases and variant calling from same-individual RNA and DNA-seq experiments from different species, allowed us to predict RNA-editing events using both primary sequence and secondary structure. Then, we devised a method for assessing conservation or divergence in the molecular mechanisms of editing completely *in silico*: the cross-training analysis. This novel method not only helps to understand the conservation of the editing mechanism through evolution but could set the basis for understanding how it is involved in several human diseases.

## Introduction

Gene regulation is without a doubt the Rosetta Stone of Genetics in the XXI Century. Among the different post-transcriptional modifications behind gene regulation, RNA editing is the one that has received less attention until a few years ago. This is especially true when compared with others as profoundly studied as alternative splicing (Hamar & Varga, 2023; Irimia et al., 2011; Irimia & Blencowe, 2012; Liscovitch-Brauer et al., 2017; Roundtree et al., 2017). In a common RNA-editing event, a single nucleotide from an RNA molecule undergoes a chemical change, turning into a different nucleotide, usually before the RNA molecule undergoes any kind of splicing (Rodriguez et al., 2012; Ryman et al., 2007). This process is present in all eukaryotic organisms (H. Liu et al., 2016; Porath et al., 2017; Takenaka et al., 2013), being the Adenine-to-Inosine (A-to-I) editing mediated by proteins of the ADAR family the most common in metazoans (Grice & Degnan, 2015; Jin et al., 2009; Savva et al., 2012). The ADAR family includes three paralog groups in vertebrates: ADAR, ADARB1 and ADARB2. ADARB1 and ADARB2 are the result of the two whole genome duplications (WGD) that took place at the origin of vertebrates, with the basal-branching cephalochordate *Branchiostoma lanceolatum* presenting only two of the three commonly found paralogs (Zawisza-Álvarez et al., 2020). Interestingly, out of the expected up to four vertebrate paralogs for each amphioxus gene, only one duplication event was conserved in the vertebrate lineage with one of the two paralogs, ADARB2, being enzymatically inactive and acting just as a binding competitor (Melcher et al., 1996; Y. Wang et al., 2019).

The ADAR-mediated A-to-I modification, although apparently small, with just one nucleotide change, can have various translational consequences. If the editing event takes place inside the coding sequence (CDS), it can have an impact on the final protein as the new inosine will behave like a guanine in any base-pairing process such as translation (Nishikura, 2016). This not only means that an amino acid can change completely but also that stop codons can be added or ignored, while different codon availability can also affect the translation. Even when happening in intronic regions, editing can have a great impact, as cryptic splice sites can arise, or modulate the specificity of microRNA targets (Nishikura, 2016).

Furthermore, A-to-I editing has been described in a myriad of processes. A prominent example is the regulation of the innate immunity in humans modulating the antiviral response. ADAR can edit the viral dsRNA thus inactivating the virus (Doria et al., 2009; Lamers et al., 2019), but this process can also dampen the interferon response (due to mismatches in the dsRNA sequence) turning ADAR into a pro-viral agent. In mice, null embryos die before birth due to stress-induced apoptosis, while ADARB1 null embryos will die young due to seizure-related complications. Alterations of the levels of editing have also been found in various diseases, such as Prader-Willi’s syndrome (Kishore & Stamm, 2006) or Alzheimer disease (Gaisler-Salomon et al., 2014). Editing in transcripts such as *GLI1* (Shimokawa et al., 2013), *AZIN1* (L. Chen et al., 2013), or *ARGHAP26* (Q. Wang et al., 2013) (in this last case the editing happening on a target of miRNAs) have been shown to be relevant in some cancers. The most prominently studied cases of A-to-I editing are those that affect the brain and nervous system of mammals and other vertebrates. Specifically, there are editing targets in key mediators of the synaptic transmission of neuronal signals, like the *GluA2, GluA3* and *GluA4* subunits of the AMPA Glutathione receptor (Lomeli et al., 1994), the *GluK1* and *GluK2* kainate-glutamate receptor (Egebjerg & Heinemann, 1993; Köhler et al., 1993; Sommer et al., 1991) or the *Nova1* splicing factor (Irimia et al., 2012). The editing in *Nova1* is a particularly noteworthy case. Nova1 is a splicing factor that regulates more than 700 splicing events, including splicing in important synaptic proteins. A specific nucleotide is a target of a conserved editing event that creates a serine-to-glycine substitution, which significantly increases the stability of the Nova1 protein. This editing event is dynamically regulated during brain development. The comparison of the editing levels in different regions of the brain shows differences in editing regulation: in *Mus musculus* there are significant regional differences in editing level, while in *Gallus gallus* all the regions have editing levels close to 100 percent (Irimia et al., 2012), suggesting that Nova-editing could have been involved in the evolution of particular regions of the mammalian brain.

Being a process as versatile and crucial as it is (Guallar et al., 2020; Higuchi et al., 2000; Lamers et al., 2019; Tonkin et al., 2002; Q. Wang et al., 2004), whether RNA editing has shaped evolution is of great interest. However, even with the several attempts made in recent years to shed light on the evolution of this process, how editing has shaped evolution is yet to be discovered (Grice & Degnan, 2015; Irimia et al., 2012; Jin et al., 2009; Takenaka et al., 2013; Yablonovitch et al., 2017; Zawisza-Álvarez et al., 2020; Zhang et al., 2023). This is mostly due to the difficulty of predicting *de novo* RNA-editing events. Little is known about the ADAR target selection mechanism besides it having to be in a dsRNA region of at least 20bp of extent (Thomas & Beal, 2017). It seems that secondary structure may have a great role in impeding or facilitating the action of ADAR proteins. This is more evident when looking at the high levels of editing of the adenosines in perfect dsRNA molecules in vitro (Thomas & Beal, 2017). These perfect dsRNA molecules have a very straightforward secondary structure which would allow ADAR to edit every single adenosine. Some studies also suggest that a complementary sequence residing in an intron that could generate a double strand in the neighbouring area of the editing site could also be necessary during editing (Wulff & Nishikura, 2010). As the target sequences or structures harbouring an adenosine that has the potential to be edited are not yet clear enough, we must rely on empirical evidence for any kind of evolutionary analysis of RNA editing. This evidence comes in the form of variant calling using same-individual genomic and transcriptomic data in order to avoid polymorphisms (Z. Wang et al., 2016), or in the form of amplification-free techniques such as Nanopore sequencing, which can identify inosines natively (L. Chen et al., 2023). Even having this empirical data and checking the conservation of the primary sequence between distinct clades, however, we can not fully ensure that mechanism is fully conserved independently from their targets.

Here we present a new approach to assess the functional conservation of the targeting mechanism independently of the conservation of editing sites using two machine learning algorithms, random forest (RF) and bidirectional long short-term memory (biLSTM) neural networks with an attention layer. RF is an ensemble method that allows building a classifier based on expert descriptors and therefore with high interpretability. On the contrary, biLSTM networks facilitate a direct approximation from sequence windows although interpretation may not be immediate. Using available RNA-editing databases and variant calling from same-individual data from different species, we trained an algorithm to predict RNA-editing events using secondary structure and primary sequence data in a species. With this, we predicted the events from different species to assess if the target selection mechanism is conserved between the two species, or whether, although sharing a similar active domain, the ADAR mechanism changed between those species. This novel method permits approaching the, until now elusive, understanding of the editing mechanisms through evolution.

## Results

### RF and biLSTM algorithms in RNA-editing prediction

The use of a random forest approach (see methods) gave us the opportunity to explore the descriptors that are most used to determine the potentiality of an RNA sequence to be edited. We ran four analyses with the RF algorithm, using a local window (Fig. 1 A) of 50, 200, 500, and 1000 nucleotides. All performed similarly well, reaching an accuracy above 75% (Supp. Fig 1). Nevertheless, there are some changes in the traditional descriptors depending on the window size used especially between 50 and 1000nt windows. Notably, the “global double strand maximum size” descriptor is highly used in both cases (Fig. 1 B).

**Figure 1.**
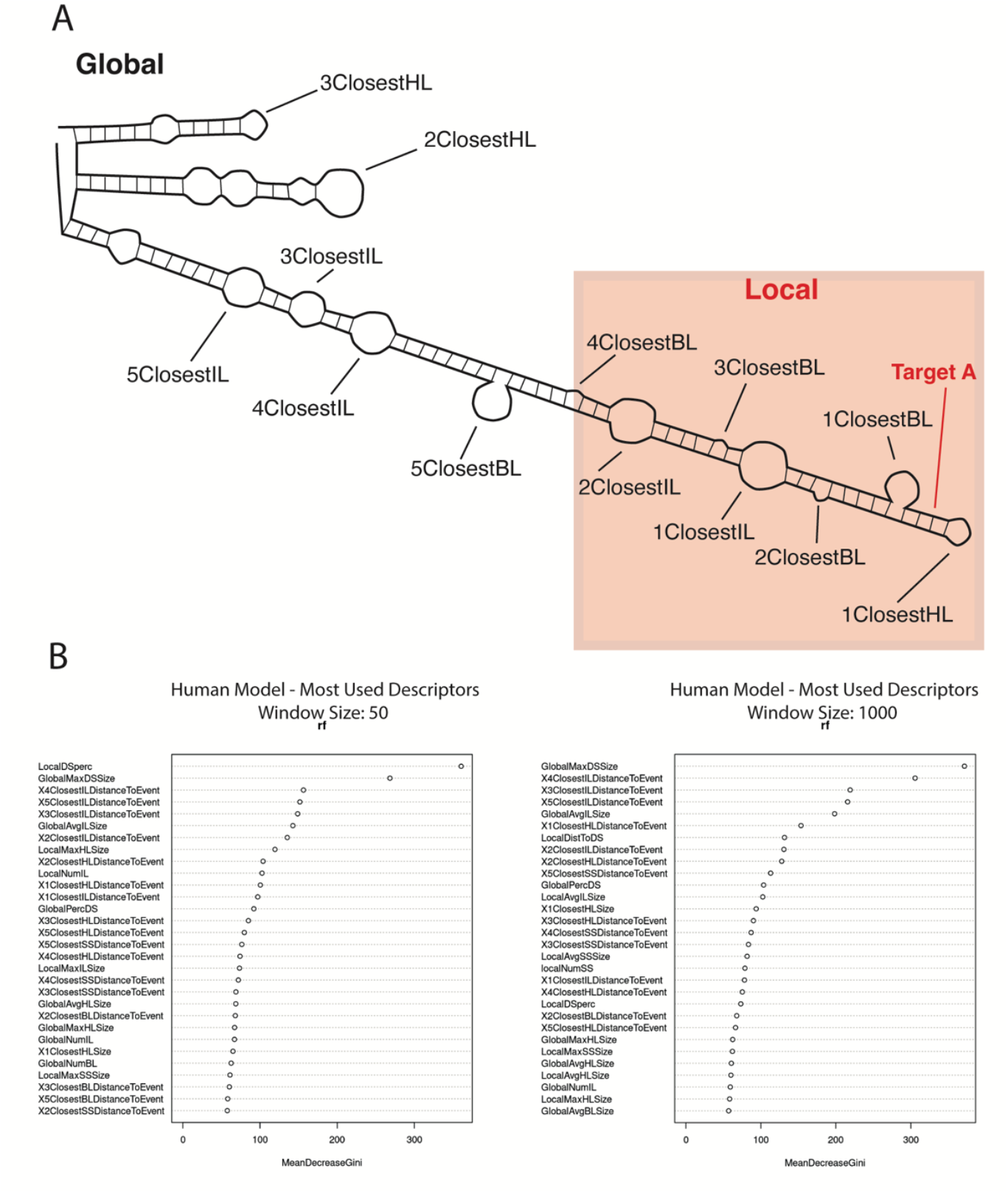
Random forest global and local results. **(A)** Schematic representation of an RNA molecule with some of the structures used as descriptors. The local window (red square) provides the data for the local descriptors, while the global descriptors use the whole molecule. Target adenosine tagged in red. the Xclosest descriptors refer to the Xth feature of that type closest to the target adenosine, independently of the local window **(B)** List of the most used descriptors in the RF analisis for the 50 and 1000 nt local windows using human data. See Supp. Methods Table 1 for the complete descriptor dataset and their definition.

On the other hand, we used a biLSTM with an attention layer (see methods; Fig. 2) with two channels, one for the pre-mRNA sequence and another one for the predicted secondary structure. Using a sliding window of 50+1+50 nucleotides, we obtained an accuracy of almost 95% using balanced datasets (Fig. 3 A). We also trained again the model using each of the two channels separately to see how they affect the ability to predict. This way, the accuracy obtained by the trained model changes when just the secondary structure channel was used (84,6%) but when using just the sequence channel it remains similar to using both channels (94,7%) (Fig. 3 B). If we explore the similarities in sequence and structure of the positive cases, we cannot see any distinguishable pattern (Fig. 3 C).

**Figure 2.**
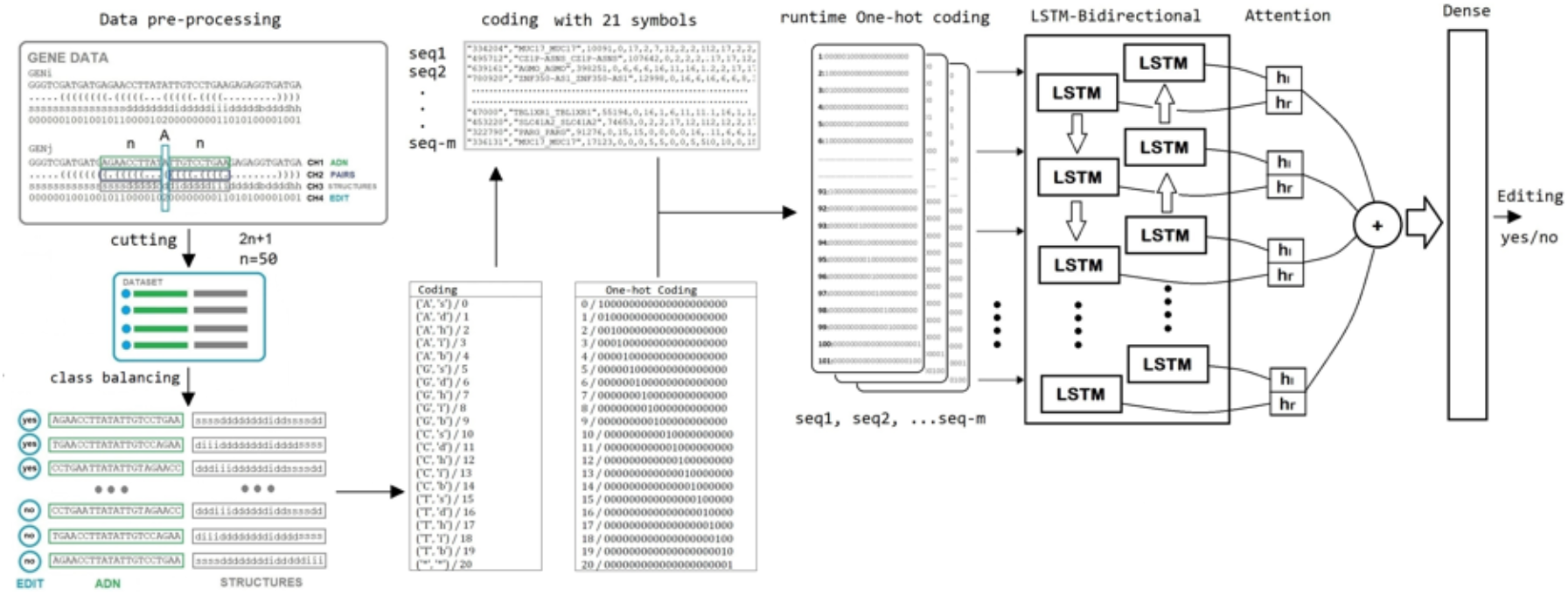
Diagram of the biLSTM data flow. From the raw data to the editability decision output. See Supp. Methods.

**Figure 3.**
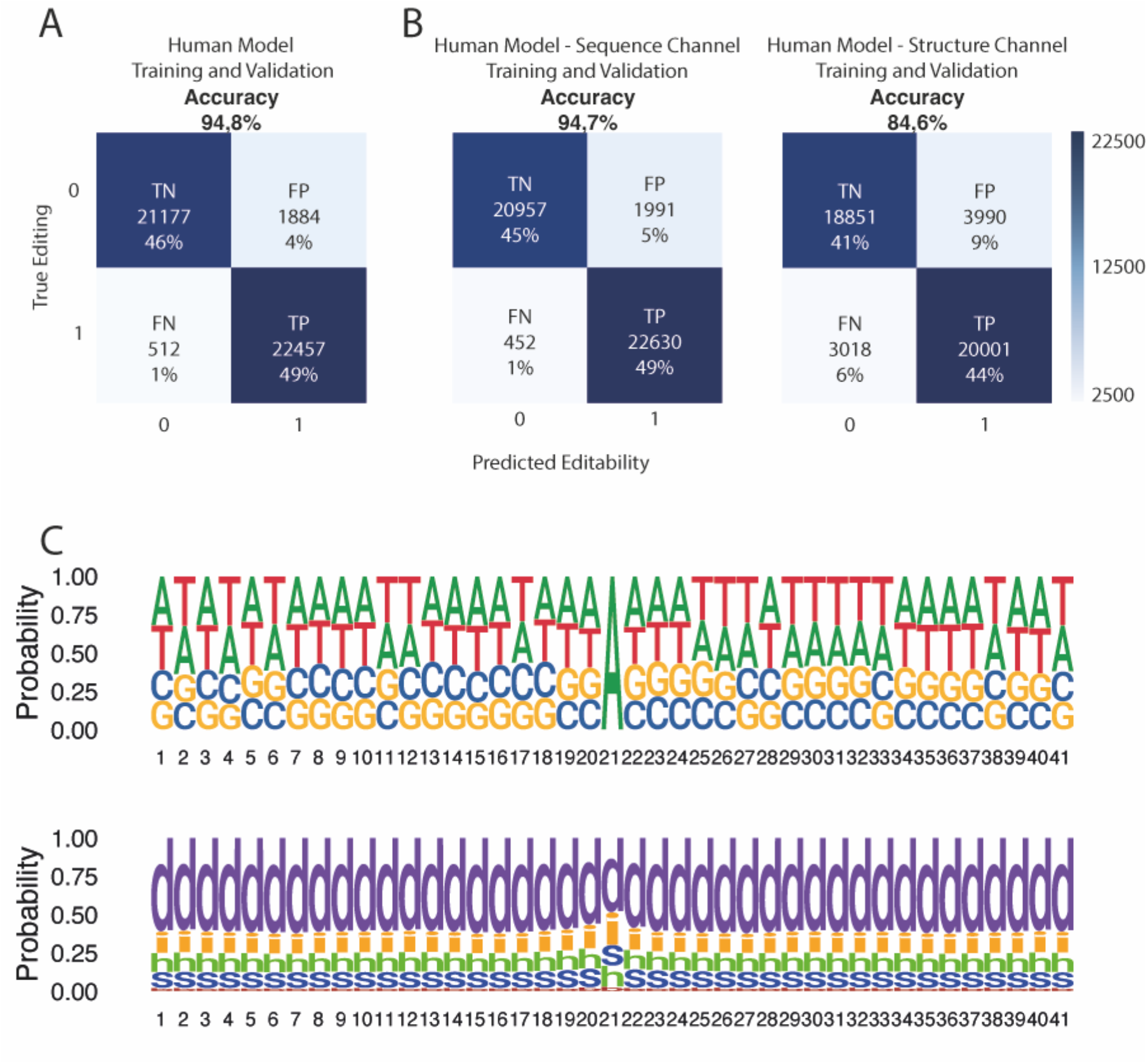
Sequence and Structure channels in DL using human data. Confusion matrices for DL analysis of sequence and structure channels from human dataset combined **(A)** or as single-channel **(B)**. True negative (TN), true positive (TP), false negative (FN) and false positive (FP) percentages have been rounded. **(C)** True positive logos for a 20+1+20 window for sequence and structure. Nucleotide 21 is the editable adenine.

### Benchmarking the algorithms with previous RNA-editing prediction attempts based on machine learning

With the human dataset predictions accuracies, we can now assess how our algorithms perform versus already existing data of machine learning predictions obtained from the bibliography (R. Chen et al., 2023; W. Chen et al., 2016; K. Liu & Chen, 2020; Xiao et al., 2018). As we can see, although our RF algorithm does not rank near other available methods, our biLSTM algorithm is the best-performing one as well as the only one using the full extent of the REDIportal database.

### Predicting a dataset using editing proportions as in a case of *de novo* prediction of RNA-editing events

Changing the balanced dataset for a dataset more akin to what we can find in a real-case scenario, gives us clues on how our prediction algorithm would perform when used for predicting new RNA-editing events. Using the full sequences of 10 random genes (as well as 20 and 30, see Supp. Figure 2), we ensure a proper data set with editing frequencies similar to the ones present in nature to benchmark our trained model. Interestingly, although the accuracy when predicting is just below 95%, the highly unbalanced nature of the dataset results in the number of false positives greatly surpassing the number of true positives (Fig. 4 A). If we explore the internal score distribution, there is a slight difference between true and false positives, and true and false negatives (Fig. 4 B).

**Figure 4.**
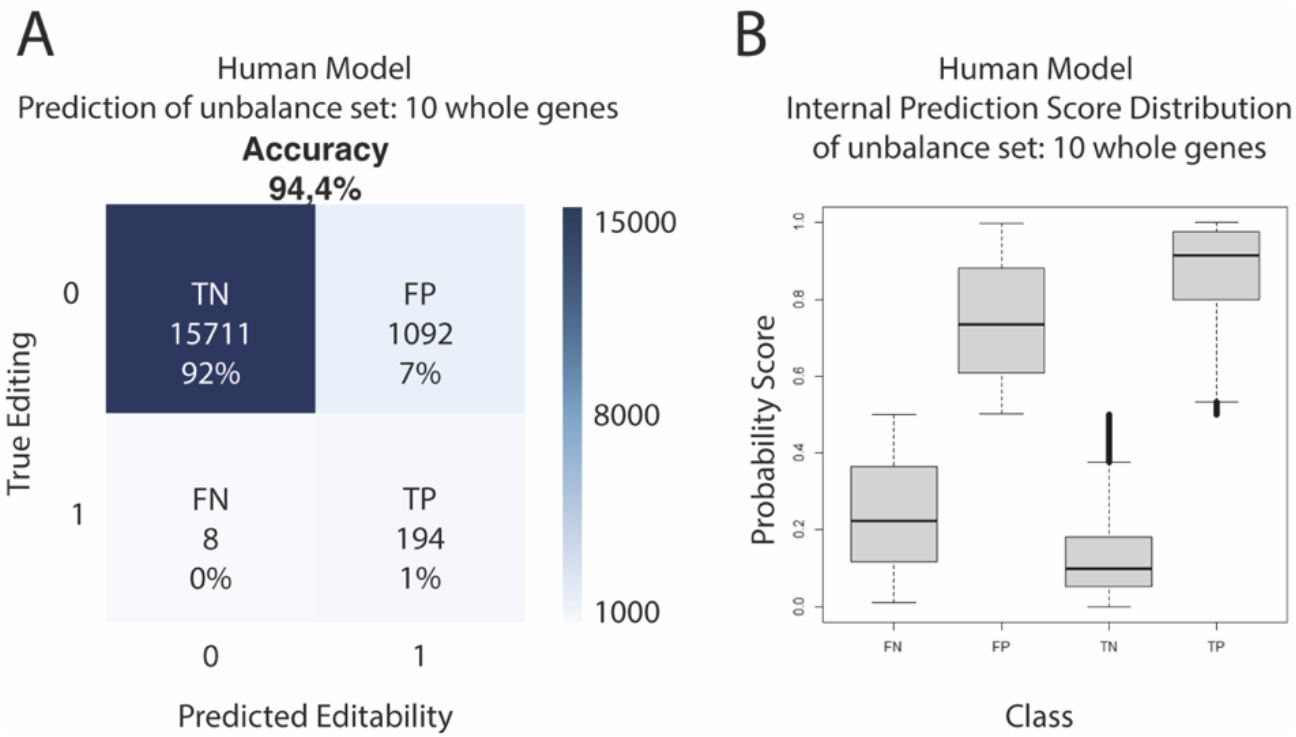
Prediction of an unbalanced dataset using DL model. **(A)** Confusion matrix for the prediction of editability in 10 whole human genes using DL model. True negative (TN), true positive (TP), false negative (FN) and false positive (FP) percentages have been rounded. **(B)** Boxplot of the internal prediction of editability score distribution for 10 whole human genes for True negative (TN), true positive (TP), false negative (FN) and false positive (FP).

### biLSTM training and predictions on non-human data

To further understand the RNA-editing process we trained the model using two additional datasets, one from a mammal (*Mus musculus*) and another one from a teleost (*Trachurus trachurus*). The mouse dataset came from the same database as the human dataset, albeit with fewer annotated RNA-editing events, while the mackerel dataset was obtained from the same individual RNA and DNA, thus being a narrow snapshot of the editome at the moment of collection. In both cases the accuracy is lower than the obtained using human data with a slight bias towards declaring an adenosine as non-editable (Fig. 5). This is especially true when using the mackerel dataset, with a 73,4% accuracy and almost 18% RNA-editing events being flagged as non-editable adenosines (Fig. 5 B).

**Figure 5.**
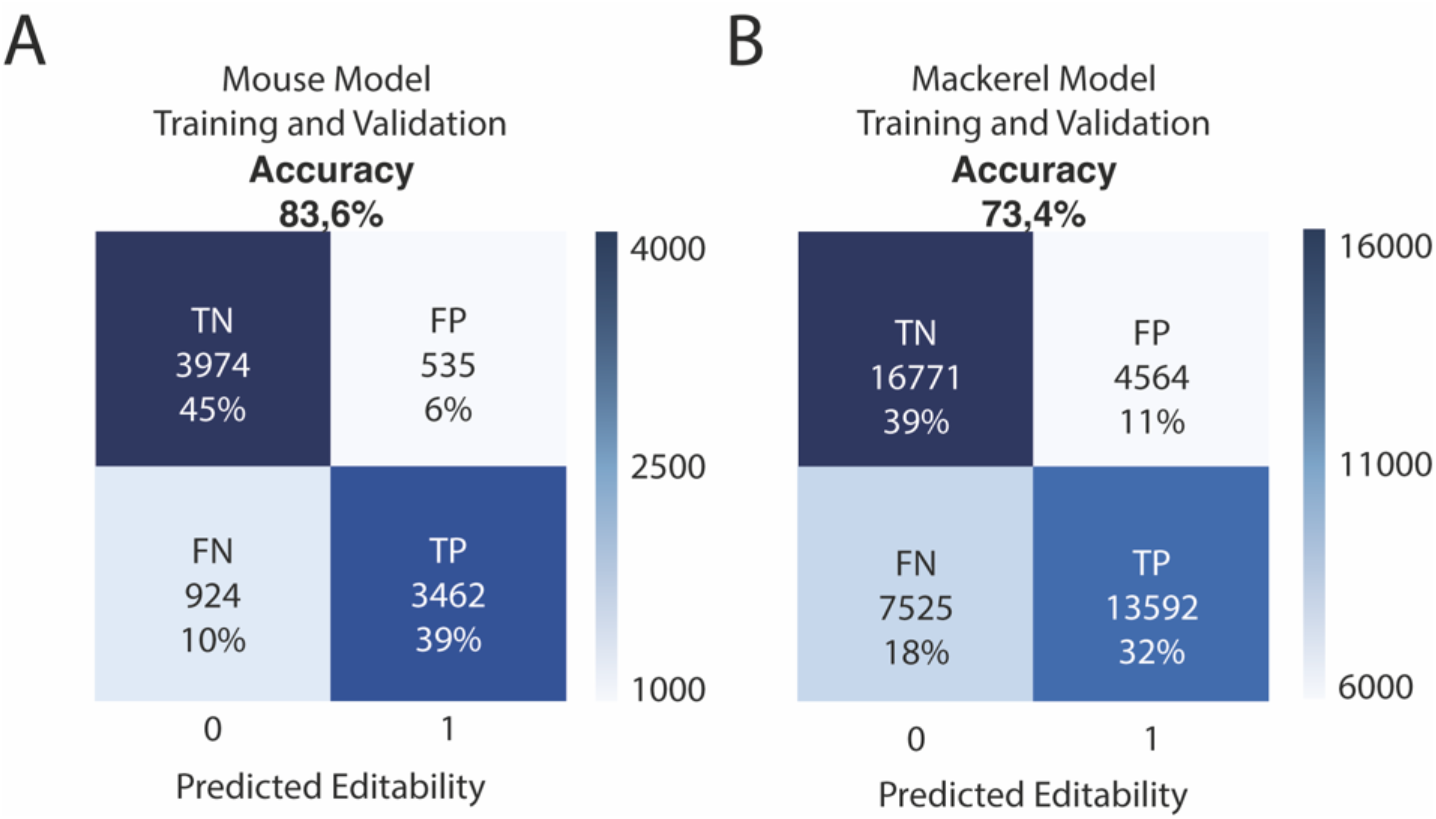
Mouse and mackerel DL analysis. Confusion matrices for models generated using mouse data **(A)** or mackerel data **(B)**. True negative (TN), true positive (TP), false negative (FN) and false positive (FP) percentages have been rounded.

### Cross-training as a tool to infer mechanism conservation *in silico*

With the data available, we explored how well an algorithm trained with the data of one species performs in predicting the data of the other species (Supp. Fig. 3, Table 1). The cross-training shows that, when trained on mammal datasets, each model can predict the other mammal’s dataset with better accuracy than the baseline of a blind random prediction (50%). When predicting *T. trachurus* data, the algorithms trained with human and mouse data achieved only 50% and 51%, respectively. Similarly, the algorithm trained with *T. trachurus* data achieved an accuracy of 52% predicting on the human dataset and 48% on the mouse dataset with similar results obtained using the algorithm trained with human data predicting on an octopus dataset (Supp. Fig. 4). Interestingly, although the algorithm trained on human data predicts better on human than the algorithm trained on mouse data when predicting on mouse data (95% vs 84%), the mouse-trained algorithm predicts better on human data (76%) than the other way around (63%). Analysing the distribution of the predictions, there is a clear tendency towards a negative prediction on the human-trained algorithm predicting mouse editable adenosines (Fig. 6 A).

**Figure 6.**
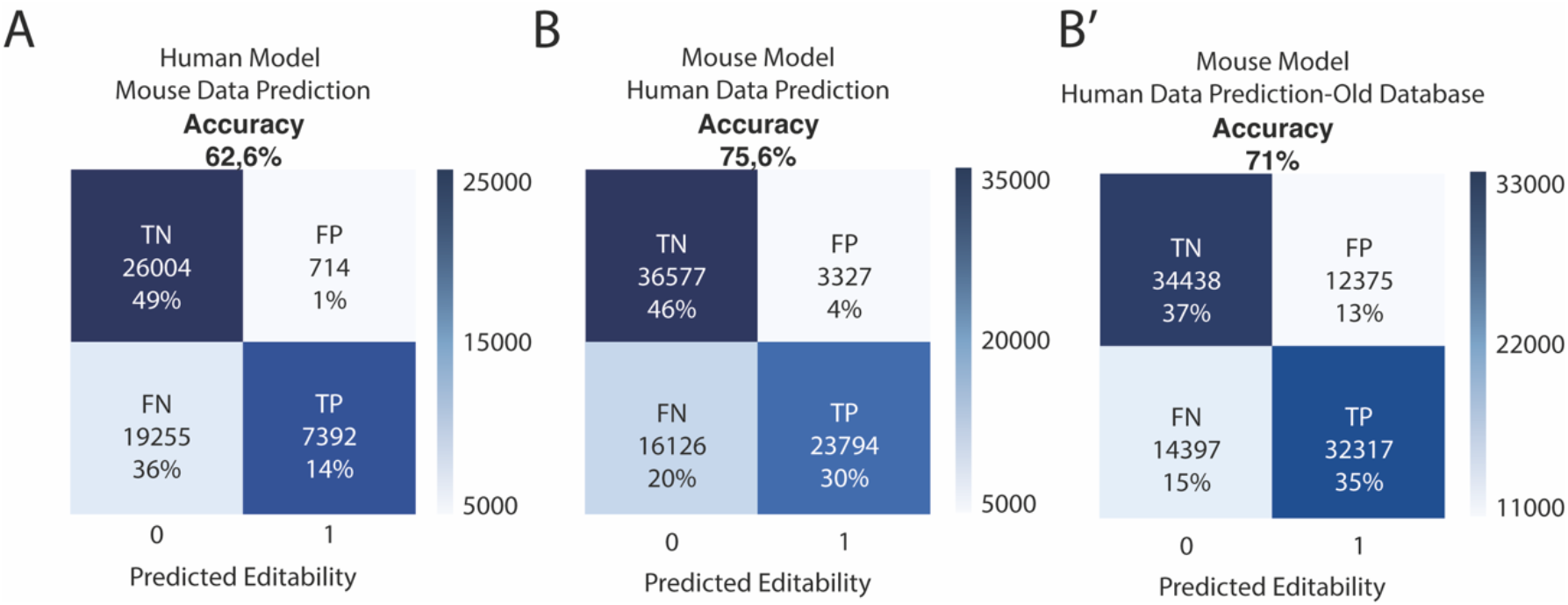
Differences of Mouse model predicting in Human new and old databases. Confusion matrices for models generated using human data predicting on mouse data **(A)** and using mouse data predicting on human data **(B)** and predicting on an old version of the used human database **(B’)**. True negative (TN), true positive (TP), false negative (FN) and false positive (FP) percentages have been rounded.

## Discussion

### Using machine learning to predict RNA-editing

#### Random Forest

Our data shows how a machine-learning approach is able to learn the RNA-editing signal. Although the random forest approach is not as accurate as the biLSTM algorithm, it is still well above the threshold expected by a random prediction (Murdoch et al., 2019) (Fig. 1). This may be due to the fact that the descriptors used in the random forest are not the most suitable for the task. Albeit they were curated by us taking into account all the previous work on secondary structure and RNA-editing (Eggington et al., 2011; Eifler et al., 2013; Thomas & Beal, 2017; Yang et al., 2006; Yeo et al., 2010), some unknown descriptors may be missing. Even so, across all the different Random Forest models there is consistency among the most frequently used descriptors: the size of the largest double-strand fragment in the whole molecule (GlobalMaxDSSize) and the distance to the target adenosine of the 4th and 5th closest inner loops (X4ClosestILDistanceToEvent and X5ClosestILDistanceToEvent) (Fig. 1 B, Supp. Fig 1). The GlobalMaxDSSize descriptor may be relevant for discriminating along the decision tree, as it is a value describing the whole RNA molecule. Any RNA molecule will have either a mixture of editable and non-editable adenosines or all non-editable adenosines. Thus, the global parameters may play a role in discriminating between these two groups (Rigatti, 2017). The relevance of the 4th and 5th closest inner loops is a bit puzzling, as it is counterintuitive that distant features are more relevant than closer ones. This could be again an early discriminating descriptor between the two aforementioned groups. The most relevant local descriptor can be found when the local window is set to 50 and 200 nucleotides (Supp. Fig 1). Here the local percentage of double-strand (localDSperc) is clearly the most used descriptor, which may mean that, with smaller local windows, the percentage of double-stranded nucleotides around the adenosine gains importance to assess the editability. Remarkably, changing the local window affects the relevance of some of the descriptors, while achieving very similar accuracies in all cases (Supp. Fig 1). This could mean that with our curated descriptors there are several ways to predict RNA editing. In the end, we see that other than the first two or three descriptors for each model, the frequency of the other descriptors remains similar, which supports the idea of a very complex decision-making process using a high number of different input variables. Although promising, our RF algorithm falls short of achieving the accuracies observed using other machine-learning methods for RNA-editing prediction (Table 1). This could be due either to the RF algorithm used or, most probably, to the curated descriptors selected being based solely on secondary structure information obtained from Linearfold.

**Table 1.**
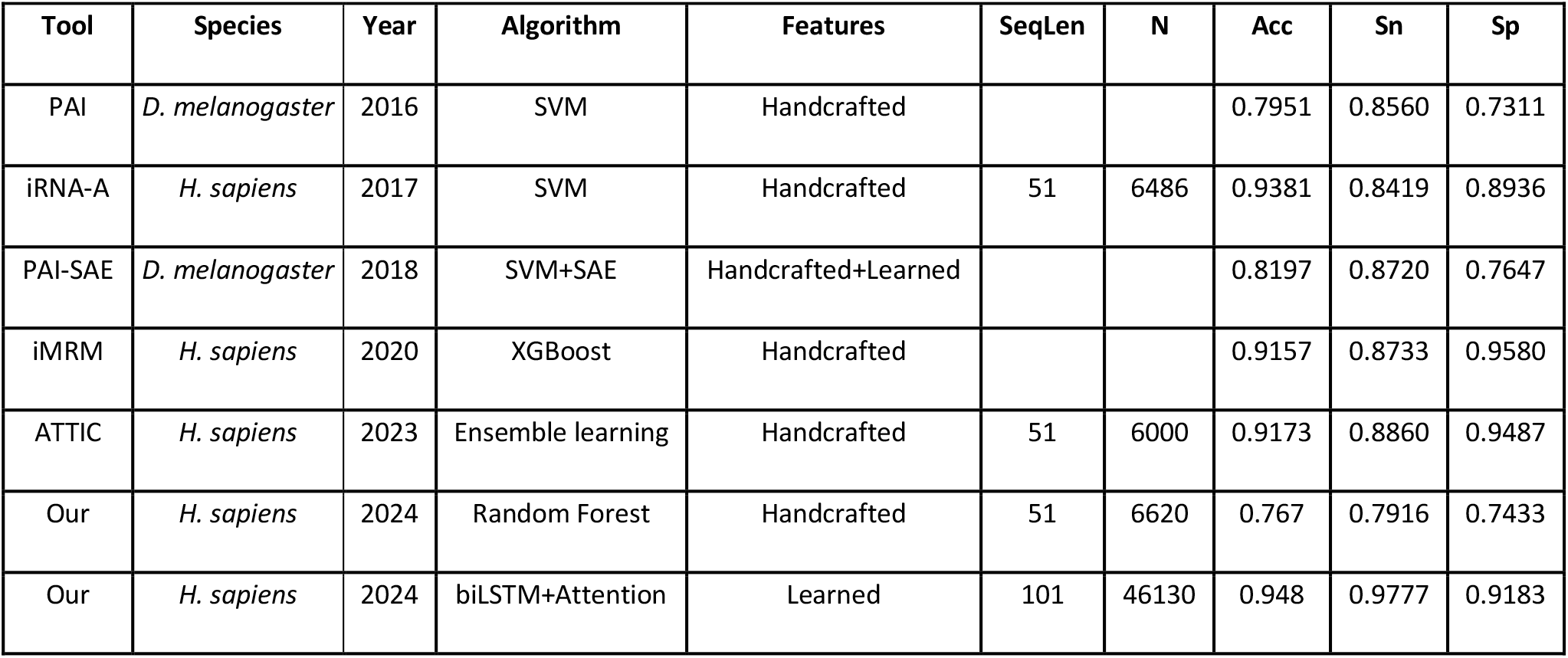
Accuracies of already existing machine learning methods found in the bibliography. SeqLen: Sequence Length analysed; N: Number of sequences used for training; Acc: Accuracy achieved; Sn: Sensitivity achieved; Sp: Specificity achieved.

| Tool | Species | Year | Algorithm | Features | SeqLen | N | Acc | Sn | Sp |
| --- | --- | --- | --- | --- | --- | --- | --- | --- | --- |
| PAI | *D. melanogaster* | 2016 | SVM | Handcrafted |  |  | 0.7951 | 0.8560 | 0.7311 |
| iRNA-A | *H. sapiens* | 2017 | SVM | Handcrafted | 51 | 6486 | 0.9381 | 0.8419 | 0.8936 |
| PAI-SAE | *D. melanogaster* | 2018 | SVM+SAE | Handcrafted+Learned |  |  | 0.8197 | 0.8720 | 0.7647 |
| iMRM | *H. sapiens* | 2020 | XGBoost | Handcrafted |  |  | 0.9157 | 0.8733 | 0.9580 |
| ATTIC | *H. sapiens* | 2023 | Ensemble learning | Handcrafted | 51 | 6000 | 0.9173 | 0.8860 | 0.9487 |
| Our | *H. sapiens* | 2024 | Random Forest | Handcrafted | 51 | 6620 | 0.767 | 0.7916 | 0.7433 |
| Our | *H. sapiens* | 2024 | biLSTM+Attention | Learned | 101 | 46130 | 0.948 | 0.9777 | 0.9183 |

**Table 1a.**
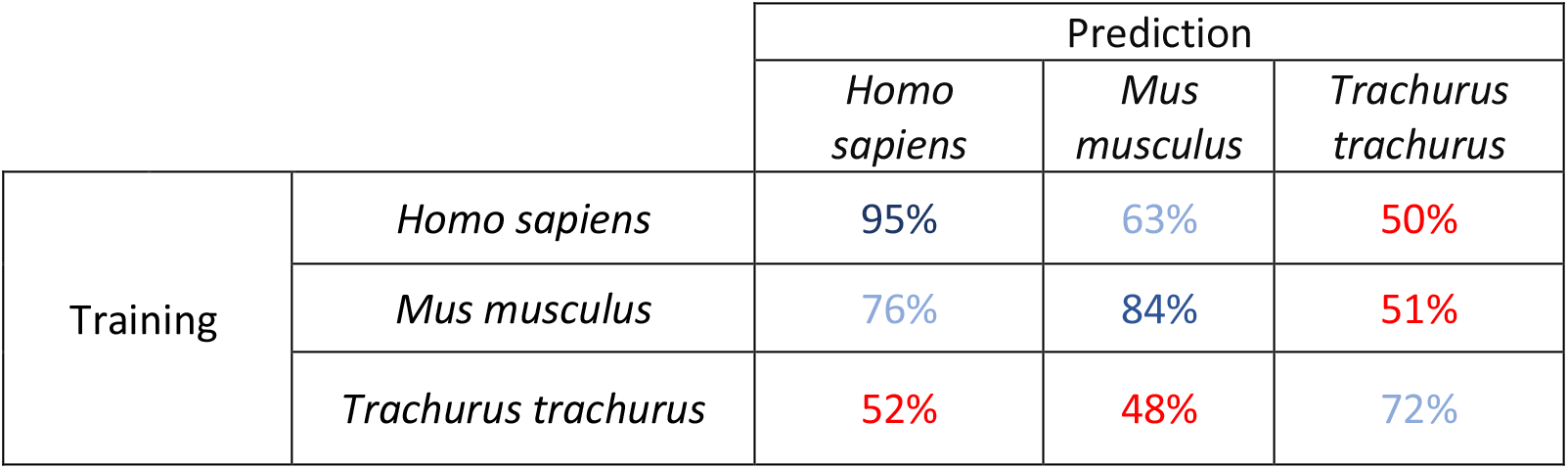
Cross-training accuracies of each possible pair of the three species analysed.

#### biLSTM algorithm

Looking closely at the biLSTM predictions, we can see that the accuracy using both (sequence and structure) channels is almost the same as the accuracy using just the sequence channel (Fig. 3 B). But even with just the structure channel, accuracy is still well over 80%. These biLSTM input single-channel experiments allow us to infer that the secondary structure is a key element for discriminating between editing and non-editing, as suggested in previous works (Thomas & Beal, 2017). However, the lower accuracy when predicting with the structure channel may mean that the biLSTM algorithm is better at predicting secondary structures from the 101-nucleotide window of the sequence channel than the specialised software (Huang et al., 2019) from the complete molecule. Another option could be that using the structure channel narrows all the possible structures to one, as this channel eliminates the sequence information (Huang et al., 2019; Mathews et al., 2010). Meanwhile, using the sequence channel would allow all the possible secondary structures to be predicted. In addition, the apparent lack of enrichment or motive seen in the separate logos for sequence and structure (Fig. 3 C) must be due to the existence of multiple highly different signals that allow ADAR to access the editable adenosine, which could imply that not all the edited adenosines have a 3’ enrichment of guanosines. Although this implication seems to conflict with the results obtained in other works (Zhang et al., 2023), it could be simply the use of a more diverse dataset, or the fact that in this study we did not differentiate between ADAR editing or ADARB1 editing. Remarkably, we can see how when compared to the existing methods (Table 1) our biLSTM algorithm performs the best in terms of accuracy using human data, even taking into account the huge differences in the training datasets, with our dataset being the whole A-to-I editing events REDIportal database.

With the data presented here, not only can we consolidate the role of the secondary structure in the RNA-editing target-selection mechanism, but we can also narrow the spatial window of the mechanism down to ±50 nucleotides from the edited Adenosine (Wulff & Nishikura, 2010; Zhang et al., 2023). This is certainly true in the two species where we have successfully learned to predict with >80% of accuracy, *Homo sapiens* and *Mus musculus*. Regarding the *de novo* prediction capabilities of both algorithms, neither is accurate enough to compensate for the huge disproportion of edited vs non-edited adenosines. One possible way of discriminating between the true and false positives could be to consider their prediction score, hindering the sensitivity of the prediction in the process. While some of the false positives detected in the unbalanced dataset could indeed be non-described RNA-editing events, it remains difficult to differentiate them from the actual (and quite more frequent) false positives. But if we compare our results from human data with other analyses using experimental RNA-editing evidence from 10 human transcripts obtained from nanopore (L. Chen et al., 2023) we see similar levels of edited sites. In addition, Chen et al. reported only a difference of 8 edited sites from the REDIportal database.

### Differences in accuracy between human and non-human data

Although *Mus musculus* and *Trachurus trachurus* prediction accuracies are significantly above random chance, they are 10 and 20%, respectively, below the accuracy obtained in *Homo sapiens*. These differences could be explained by the different characteristics of the datasets, for example, the different sizes. However, the results obtained adjusting the training datasets to the same number of events available for *Mus musculus* yielded very similar results in *Homo sapiens* (Supp. Fig. 5). This means that between those two species, the different accuracies do not arise from the number of events but from the kind of events available in each dataset. In this light, we decided to train the model with an older version from the human database as, akin to the mouse dataset, it will have a more generic set of RNA-editing events than its newer counterpart (Supp. Fig 6) (Mansi et al., 2020; Picardi et al., 2017). The algorithm trained with the older version of the database, nonetheless, performed similarly to the newer one, with a slight decrease in accuracy, meaning that even with the same number of entries, the more uncommon editing examples available, the more accurate the predictions become. As for *Trachurus trachurus*, we found a severe decrease in accuracy when adjusting its dataset size to the *Mus musculus* dataset. This could be caused by the lower quality of the dataset, as its origin is a single RNAseq experiment (coupled with same-individual genomic data) from the Darwin Tree of Life project (Genner et al., 2022). Nonetheless, the possibility of some of the false positives being non-described edited sites was also considered. However, due to the scarceness of editable sites when compared with non-editable sites, and the samples being balanced on editable sites, it should not impact the overall accuracy of the analysis.

### Cross-training and mechanism conservation

One of the most promising applications derived from the machine-learning approach studied here is the inference of functional conservation completely *in silico*. With this in mind, we used cross-training: training with datasets from one species and testing on datasets from other species. If the mechanism is fully conserved between two species (that is, the patterns that ADAR recognizes are the same) with a similar completeness database, the accuracy between their cross-trainings should be similar. Here we see how this happens between the cross-trainings from the old and the new human databases (Mansi et al., 2020; Picardi et al., 2017) (Supp. Fig. 6). In the case of the cross-trainings between *Homo sapiens* and *Mus musculus*, we show how, although similar, the mouse-trained algorithm performs better on the human dataset than the other way around. The reason for this may be a minor functional difference coupled with the aforementioned lower completeness of the database from *Mus musculus*. From the bias towards false negatives present in the human-trained algorithm, it may be inferred that mouse-specific structures are being misclassified as non-editable. This would explain the higher balance between false negatives and false positives when predicting the human dataset with a mouse-trained algorithm as well as the already mentioned bias and the different accuracies (Fig. 6). These differences are more obvious when cross-training the old human database with *Mus musculus* (Fig. 6 B’) and are also observed in the Random Forest cross-training tests (Supp. Fig. 7).

For *Trachurus trachurus* cross-training, in all cases, the accuracy is around 50% which is expected for a random prediction (Murdoch et al., 2019). While the accuracy of the biLSTM algorithm trained in *T. trachurus* was not as high as the one from human or mouse, we did not expect such low performance in the cross-trainings. The main reason for the inability to predict in *T. trachurus* when training the algorithm in human or mouse (or the other way around) may well lay in the differences in homeostatic temperature affecting the secondary structure of the RNA molecules (Anania, 2023) with similar results obtained in the cross-training using human data to predict octopus editable sites (Supp. Fig. 4). If we analyse the single-channel biLSTM results from *T. trachurus*, we can see how we fail to predict above random chance when using just the secondary-structure channel (Supp. Fig. 8). This could mean that here the secondary structure prediction software used (Huang et al., 2019) is not working as intended in the case of the cold-blooded mackerel, with the biLSTM algorithm completely relying on the sequence channel.

Our results demonstrate the power of machine learning approaches to predict RNA editing events. However, despite the extremely high accuracy reached, we are not yet able to use these algorithms to predict de novo editing events reliably due to the unbalanced nature of edited vs non-edited adenosines. Nonetheless, thanks to our new cross-testing approach, we can further understand the differences in RNA editing between different species, and how these differences could have shaped evolution. This opens the door to investigate whether some species have a fast-evolving RNA-editing machinery, or if the absence of one of the ADARs can reshape the RNA-editome.

## Methods

A more detailed section can be found in Extended Methods

### Origin of the RNA-editing and genomic data

We obtained the human and mouse RNA data from REDIportal (Picardi et al., 2017), as well as the RefSeq gene notation and the standard genome assemblies (hg38 for human and mm10 for mouse). We also obtained an older version of the human REDIportal database from the authors. For mackerel, we used the DNA-seq and RNA-seq data from the Darwin Tree of Life(Genner et al., 2022), which is from the same specimen. We also used the genome assembly and gene annotation from the Darwin Tree of Life. We aligned the DNA reads using Magicblast (v1.6.0)(Boratyn et al., 2019) and the RNA reads using bowtie2 (v2.4.2) (Langmead & Salzberg, 2012) and then used the SAMtools (v1.15.1)(Danecek et al., 2021) and bcftools (v1,11)(Danecek et al., 2021) libraries to obtain separately the DNA and RNA SNVs in vcf format. Then we filtered the A-to-G variants that appeared only in the RNA SNVs, filtering out the polymorphisms from the DNA SNVs. We also set a minimum depth of 10. This resulted in our accepted mackerel RNA-editing positions. A more detailed process can be found in Extended Methods.

### General pipeline for constructing the Random Forest and Neural networks datasets

We used mostly our own programs to get the datasets for both the random forest and neural networks approach. We extracted the pre-mRNA sequences that had editing events in them and predicted the secondary structure using linearfold (Huang et al., 2019). We annotated the information about the secondary structure in two different ways: for random forest we have, for each adenosine in a dataset, a series of descriptors that give information about the features of the secondary structure both close to the particular adenosine and for the whole molecule. For the neural networks approach, we have an input two-channel sequence, one channel with the pre-mRNA sequence and the other with the type of secondary structure feature each nucleotide is in. We then cut the sequences in windows of 50+1+50 nucleotides around a particular adenosine. Both for random forest and neural networks, we have positive datasets, with potentially edited adenosines, and negative datasets, with non-edited adenosines. For both approaches we divide the datasets into training and testing. For random forest we use R 4.2.1, while the neural networks approach, we used a bidirectional biLSTM neural networks model implemented in keras. A more detailed process can be found in Extended Methods.

### RF

Random Forest is a supervised ensemble algorithm based on decision trees. Each decision tree is built using a random sample of the original data (bootstrap) and a feature randomness selection, in this way the forest of uncorrelated trees is created that will serve to make a prediction by committee with better performance than if it were individual. Furthermore, it is possible to obtain which features are most relevant to build the predictor, which makes this algorithm easy to interpret. The R package randomForest has been used as the implementation.

Before running the random forest, the presence of missing values and the degree of variability of the descriptors must be checked.

The descriptors used for the random forest datasets may have missing values for some variables. XClosest descriptors may have missing data when the sequence in some type of structure is less than 5 occurrences. In this case, the sequence is removed. Another case is when local descriptors such as localAverageXXXSize there are not any occurrences XXXSize so it is not possible to compute the average. If the number of missing values is greater than 10% then the descriptor is removed and otherwise, the sequence is removed.

In addition, variables that have very little variability are eliminated since they do not provide relevant information and simplify the model.

The selection of the two hyperparameters of the random forest algorithm, number of trees and number of variables, was carried out individually for each window size 50, 200, 500 and 1000 nucleotides and organism. In all cases, it was tuned to a maximum number of 1000 trees and possible number of variables: 2, 4, 5, 7, 9, 13, 17, 19, 25, 30, 40, 50, 60, 90, 110, 120, all variables. The out of bag (OOB) score was used as a performance measure for the selection of hyperparameter values. The Gini index was used as measure of purity to create of trees.

### biLTSM

We have used a bidirectional LSTM, which processes sequences in the two possible directions, along with an additional self-attention layer. An LSTM network, long short-term memory (Hochreiter & Schmidhuber, 1997) is a type of recurrent neural network, based on a special type of recurrent unit, that solves the vanishing gradient problem present in older models. The LSTM network is capable of learning relationships, both between nearby points and between points far away into the sequence. Pre-mRNA sequences, due to their secondary structure, can present these types of spatial relationships, even between distant nucleotides in the primary sequence.

Bidirectional networks usually offer better performance than unidirectional LSTMs and also treat the tokens in a sequence in a symmetrical way.

We have also added a layer of self-attention, with the aim of trying to improve performance. The attention layer is capable of assigning different weights to different positions in each input sequence, seeking to give more relevance to the positions that are most decisive when classifying the sequence.

To develop the recurrent deep learning models we have used the LSTM implementation made by tensorflow.keras, using the layers.Bidirectional and layers.LSTM classes. A more detailed process can be found in Extended Methods.

### Software and Hardware Used

To carry out this work, the following programming environments and libraries were used:

- Python3
- Spyder IDE for Python Development
- Tensorflow 2.7.0 (Martín~Abadi et al., 2015)y Keras 2.7.0 (Chollet & others, 2015).
- Conda 4.11.0 and Anaconda Navigator 2.1.2.(“Anaconda Software Distribution,” 2020)
- JupyterLab 3.2.1. (*JupyterLab*, n.d.)
- Google Colab Pro. (*Colab.Google*, n.d.)
- The CUDA release 11.6, and cuDNN 8.3 libraries. (*CUDA Deep Neural Network (CuDNN)* | *NVIDIA Developer*, n.d.; *NVIDIA Developer Documentation*, n.d.)

We have used the following computers.

- Personal computer with GPU: We have used a laptop with Intel i7 processor, 16Gb RAM and 1TB SSD disk. In addition, it has nVidia GeForce MX450 2Gb GPU.
- Subscription to the Google Colab Pro service, which allows you to use a virtual machine with 26Gb of RAM and a Tesla T4 GPU card with 15Gb of RAM.
- AMD opteron server, Processor 6386 SE, with 500Gb of RAM, 64 processors and NVIDIA TU104GL graphics card [Quadro RTX 4000]. It was used for processing genome-wide data.

## Data availability

The authors declare that all data supporting the findings of this study are available within the paper and its supplementary information files. The software used can be found in: https://github.com/cherrera1990/RNA-editing-pred

## Acknowledgements

We would like to acknowledge Enrique Navas, Claudia Armillas and all of Jordi Garcia’s lab members for the scientific discussion. M.Z.-A. is a recipient of an FPU (Formación del Profesorado Universitario) scholarship from the Spanish Ministerio de Universidades. C.H.-U. is a recipient of a Margarita Salas postdoctoral contract from the Spanish Ministerio de Universidades. J.G-F. received funding from the Ministerio de Educación y Ciencia (grant number BFU2017-86152-P and PID2020-117820GB-I00). J.G.-F. benefit from 2021-SGR00372-GRC from AGAUR (Generalitat de Catalunya).

## Author contributions

J.G.-F., C.H.-U. and M.Z.-A. conceived the presented idea; M.Z.-A. obtained and prepared all the data used; J.P.-M. and F.R. designed and performed the DL analysis; E.V. designed and performed the RF analysis; C.H.-U. and M.Z.-A. provided the theoretical framework and contributed to the interpretation of the results. All authors contributed to the writing, provided critical feedback and helped shape the research, analysis and manuscript.

