## Supplemental Figures for "Exploring functional conservation *in silico*: a new machine learning approach to RNA-editing"

### Supplementary Results

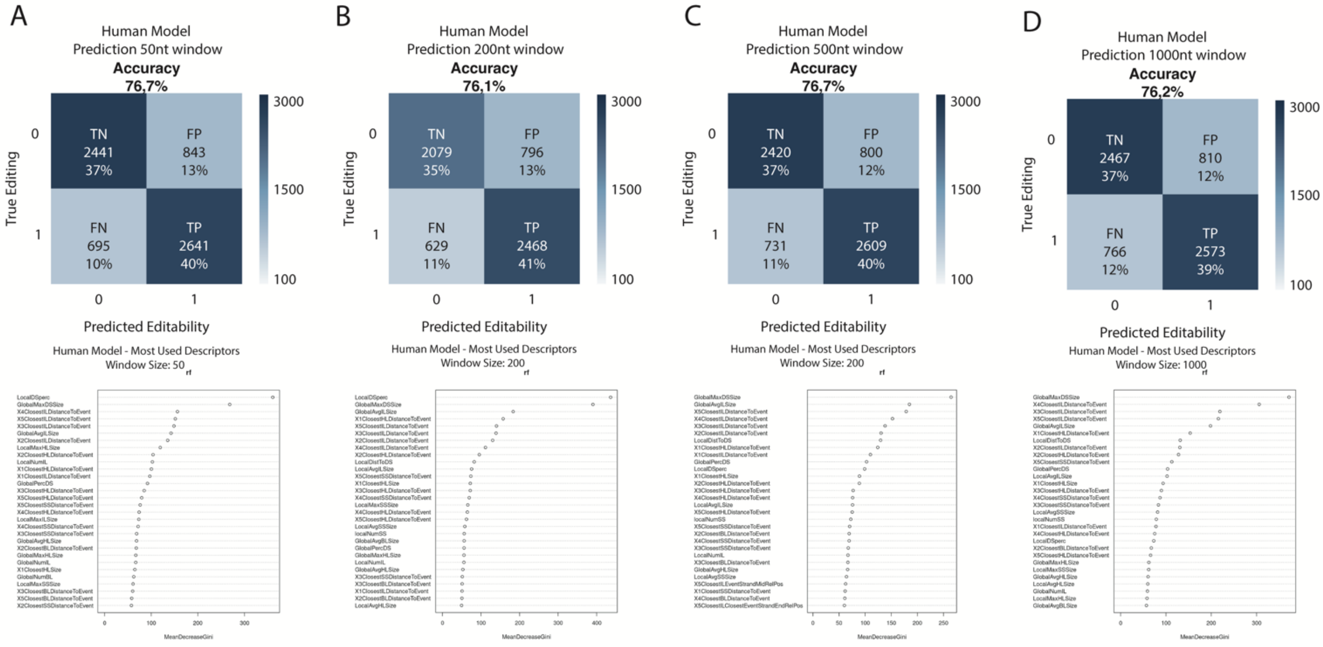


**Supplementary Figure 1. Complete RF analysis.** Confusion matrices combined with the list of the most used descriptors in the RF analysis for the 50 **(A)**, 200 **(B),** 500 **(C)**, or 1000 nt **(D)** local windows using human data. See Supp. Methods Table 1 for the complete descriptor dataset. True negative (TN), true positive (TP), false negative (FN) and false positive (FP) percentages have been rounded.

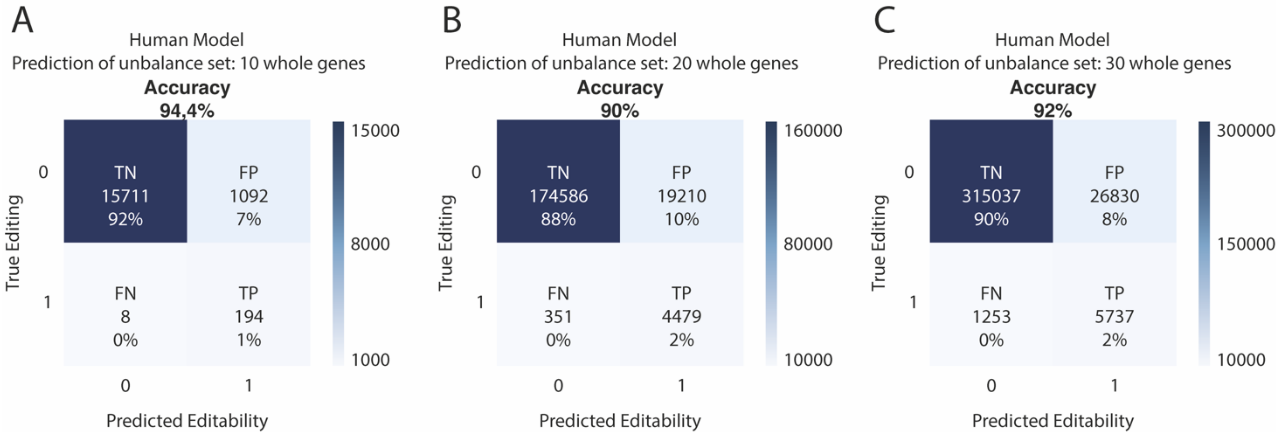


**Supplementary Figure 2. Prediction of an unbalanced dataset using DL model.** Confusion matrices for the prediction of editability in 10 **(A)**, 20 **(B)**, and 30 **(C)** whole human genes using DL model. True negative (TN), true positive (TP), false negative (FN) and false positive (FP) percentages have been rounded.

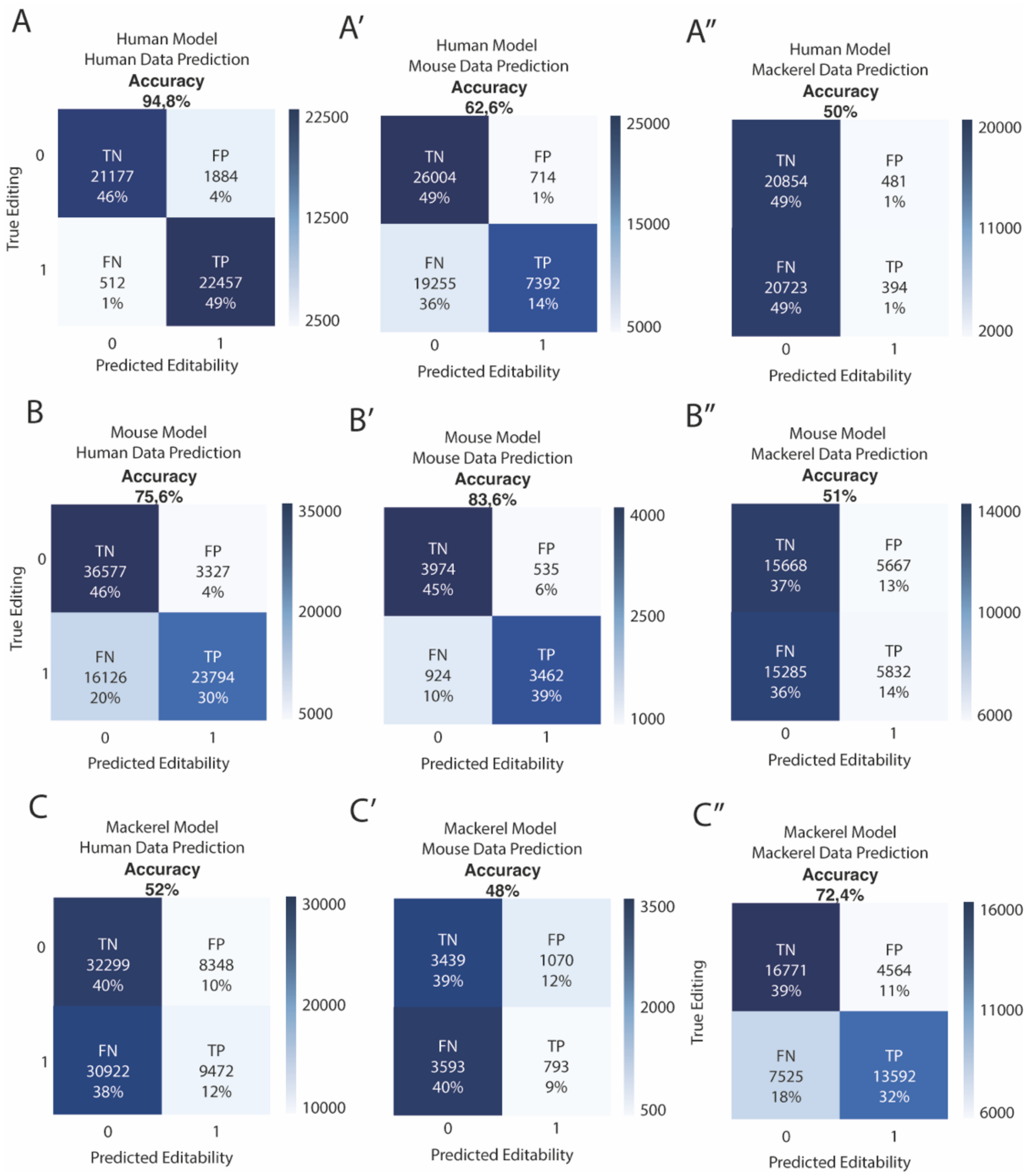


**Supplementary Figure 3. Confusion matrices for cross-training DL analysis**. Using human data predicting on human data **(A)**, predicting on mouse data **(A')** and predicting on mackerel data **(A'')**. Confusion matrices for cross-training DL analysis using mouse data predicting on human data **(B)**, predicting on mouse data **(B')** and predicting on mackerel data **(B'')**. Confusion matrices for cross-training DL analysis using mackerel data predicting on human data **(C)**, predicting on mouse data **(C')** and predicting on mackerel data **(C'')**. True negative (TN), true positive (TP), false negative (FN) and false positive (FP) percentages have been rounded.

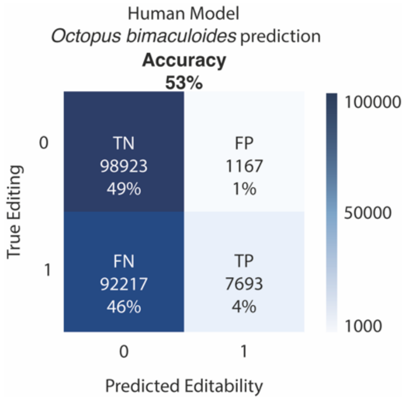


**Supplementary Figure 4. Confusion matrix for cross-training DL analysis using human data predicting on octopus data.** True negative (TN), true positive (TP), false negative (FN) and false positive (FP) percentages have been rounded.

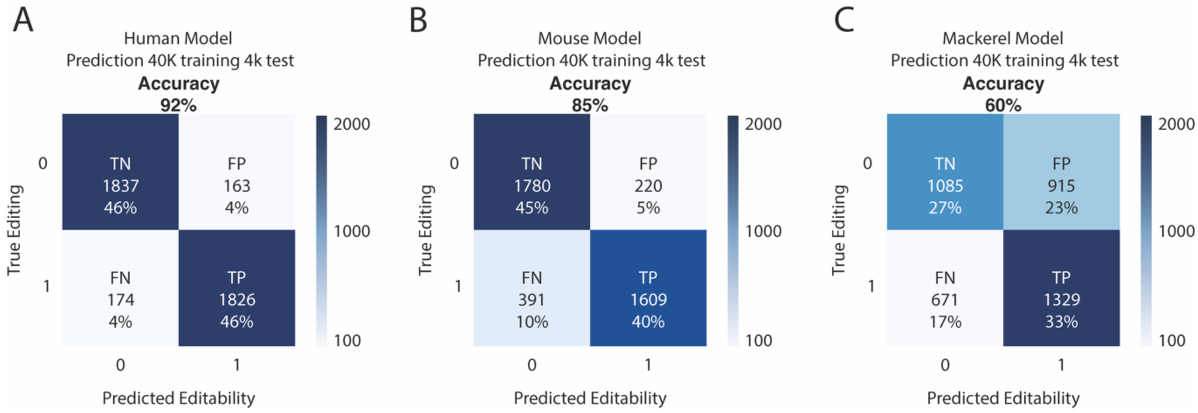


**Supplementary Figure 5. Confusion matrices for DL analysis of dataset adjusted to the same size (40k elements).** For Human **(A)**, Mouse **(B)**, and Mackerel **(C)**. True negative (TN), true positive (TP), false negative (FN) and false positive (FP) percentages have been rounded.

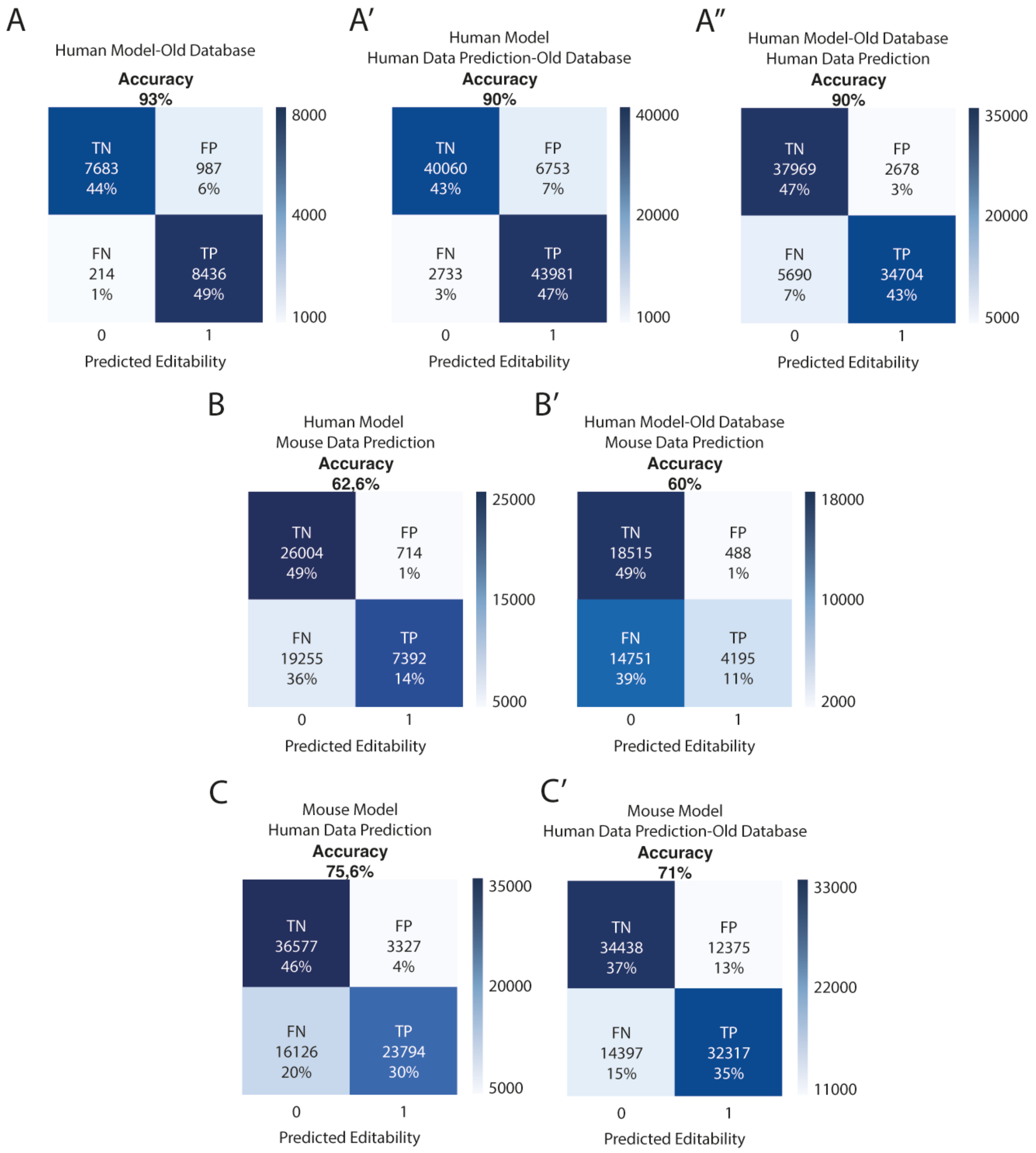


**Supplementary Figure 6. Differences of Mouse model predicting in Human new and old databases, extended.** Confusion matrices for DL analysis of Human old dataset **(A)**, DL cross-training analysis using Human data predicting on an old version of the used human database **(A')** and using an old version of the used human database predicting on human data **(A'')**. Confusion matrices for cross-training DL analysis using human data predicting on mouse data **(B)** and using an old version of the used human database predicting on mouse data **(B')**. Confusion matrices for cross-training DL analysis using mouse data predicting on human data **(C)** and using mouse data predicting on an old version of the used human database **(C')**. True negative (TN), true positive (TP), false negative (FN) and false positive (FP) percentages have been rounded.

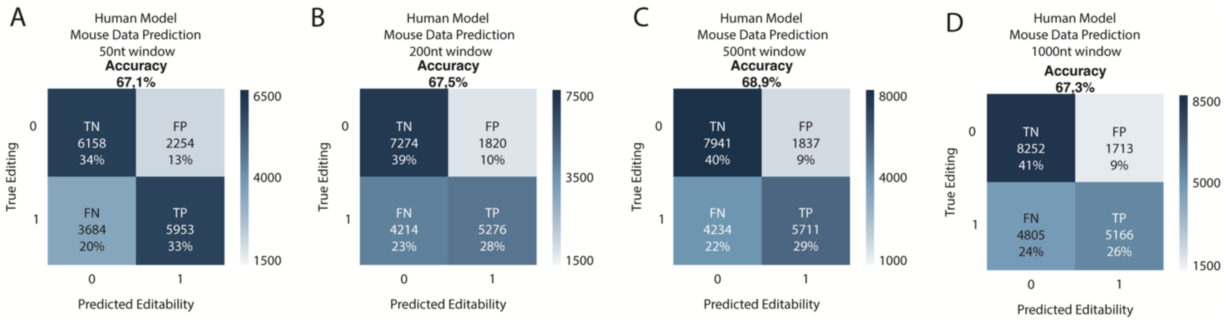


**Supplementary Figure 7. Confusion matrix for cross-training RF analysis using human data predicting on mouse data.** Using the 50 **(A)**, 200 **(B),** 500 **(C)**, or 1000 nt **(D)** local windows. True negative (TN), true positive (TP), false negative (FN) and false positive (FP) percentages have been rounded.

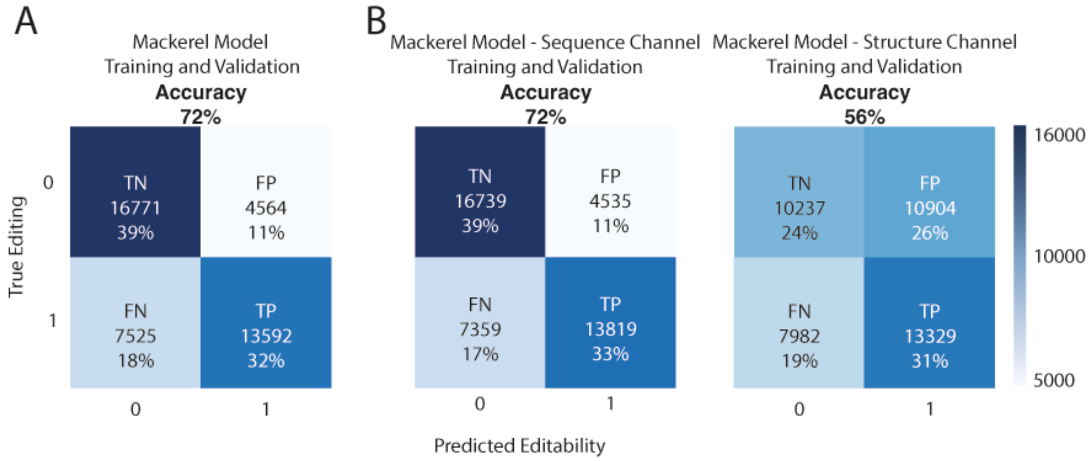


**Supplementary Figure 8. Sequence and Structure channels in DL using mackerel data.** Confusion matrices for DL analysis of sequence and structure channels from mackerel dataset combined **(A)** or as single-channel **(B)**. True negative (TN), true positive (TP), false negative (FN) and false positive (FP) percentages have been rounded.
