## Extended Methods for "Exploring functional conservation *in silico*: a new machine learning approach to RNA-editing"

### Supplementary Methods

#### Origin of the RNA-editing and genomic data

For human and mouse, we got the data from the RNA-editing database REDIportal (Picardi et al., 2017) and the standard genome assemblies (hg38 for human and mm10 for mouse) as well as the RefSeq gene notation. We also obtained an older version of the human REDIportal database from the authors.

For mackerel, we used the DNA-seq and RNA-seq data from the Darwin Tree of Life (Genner et al., 2022), which is from the same specimen. We also used the genome assembly and gene annotation from the Darwin Tree of Life. We aligned the DNA reads using Magicblast (v1.6.0) (Boratyn et al., 2019) and the RNA reads using bowtie2 (v2.4.2) (Langmead & Salzberg, 2012) and then used the SAMtools (v1.15.1) (Danecek et al., 2021) and bcftools (v1,11) (Danecek et al., 2021) libraries to obtain separately the DNA and RNA SNVs in vcf format. We used our own program, variants_filter_gff, which uses as input the vcf file and a gff file, to get the A-to-G variants that fall inside positive strand transcripts and T-to-C variants that fall inside negative strand transcripts (as well as A-to-G and T-to-C variants that do not fall inside any annotated transcript), and add the -d 10 option to filter out the variants that have a depth of less than 10. The variants_filter_gff program adds to each vcf line information about the transcripts where the variant falls, but we do not use this information later. We then used our program DNA_reads_editing_filter to filter the RNA variants with the DNA vcf file and discard those RNA variants that are detected as DNA polymorphisms. These will be our accepted RNA-editing positions in mackerel.

In order to unify the formats coming from the mackerel vcfs and those coming from the human and mouse REDIportal database, we used the linux command cut to select the relevant columns (1,2,4,5,11) from the mackerel RNA-editing file. Because variants_filter_gff can duplicate variants if they fall in more than one transcript, we later used the linux commands sort and uniq to delete repeated lines.

#### General pipeline for constructing the Random Forest and Neural networks datasets

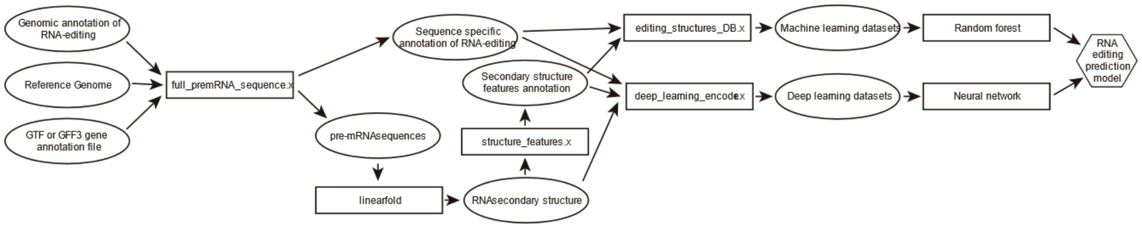


**Supplementary Methods Figure 1. General representation of the pipeline shared between the Random Forest and Neural Networks approaches.** This shows all the programs used and intermediate data generated in the steps between having the genomic positions of RNA-editing to having the particular datasets for the two approaches which will be further processed separately. Note that, for simplicity, we have omitted the non-edited adenosines datasets which undergo more or less the same processing.

We use the program locus_lengths (or locus_lengths_gff if the annotation is a gff) to get the list of the lengths of all the annotated loci (if we have individual isoforms annotated, then we take the maximum lengths, min(start) to max(end)). We use R(3.5.1) to open the loci lengths file and use the R instruction summary to get the statistical descriptors of the length. We calculate the threshold for considering a locus or gene an outlier in terms of its pre-mRNA length with the formula: Q3 + 1,5 * IQR. We use this in the next step to exclude the editing that happens in genes that are outliers in terms of length, because we found out that we get much more consistent results in the Random Forest datasets, but with the Neural networks approach there doesn't seem to be a difference between excluding outlying genes or not.

We then use the program full_premrna_sequence, using as input (Supp. Met. Fig. 1):

- either the REDIportal database lines (for human and mouse) or the correctly formatted and filtered variants file (the output from the uniq of the previous step)
- The genome sequence in fasta
- The gene annotation file (gtf or gff)

This program has two separate outputs:

- It takes the RNA-editing genomic positions annotated in the input file (only those that are A-to-G on the positive strand and T-to-C on the negative strand) and checks with the GTF or GFF file, inside of which genes falls each position. It outputs the same line as the input, but adding at the start the transformation of the genomic position to the local position within the pre-mRNA transcript of the corresponding gene. Note that with a gff file (using -g and -f options), which usually has the genes annotated directly, the program uses the gene start and end positions as the start and end of the pre-mRNA transcript. With a gtf file, which usually has only transcript annotations, the program searches for all the transcripts of each gene and considers the pre-mRNA start position as the minimum of all the starts of the transcripts of the gene and the end of the pre-mRNA as the maximum of the ends of the transcripts.
- For the genes that have at least one RNA-editing target inside its pre-mRNA, we also output the whole pre-mRNA sequence onto a separate file (there's also the option -d to divide the output file into multiple files in order to later parallelise the prediction of the secondary structure). We take the pre-mRNA sequence directly from the genome fasta file, using the pre-mRNA start and end positions as explained before.

For the full_premrna_sequence program we use the following options:

- -d 60 in order to divide the pre-mRNA sequences between 60 separate files.
- -n 20 in order to ignore any gene that has over 20% of Ns in the pre-mRNA sequence, in order to obtain good enough secondary structure predictions.
- -s with the outlier threshold we calculated using R, in order to ignore the genes that are outliers in terms of length
- - g and -f when we have gff files instead of gtf (mackerel)
- -p when in the fasta file, the header has more information than the chromosome (or scaffold) id. This separates the fasta header by whitespace and takes as the chromosome id the first part.

We then run linearfold (Huang et al., 2019) with the pre-mRNA files as input (in parallel), to get the secondary structure prediction of the sequences (Supp. Met. Fig. 1). Then we regroup all the resulting secondary structure files into a single file using the cat command. We use the program structure_features, with the secondary structures as input, in order to obtain a custom annotation of the features of the secondary structures (Supp. Met. Fig. 2). This program detects and annotates the following structures, with the annotation including the start and end positions of the feature and the sizes of the feature (for those structures where it is relevant, it includes both strands of the feature):

- Hairpin loops (HL/h)
- Inner loops (IL/i)
- Bulge loops (BL/b)
- Other single strand fragments (SSBF, SSMF, SSEF/s)
- Double strand fragments: (DS/d)

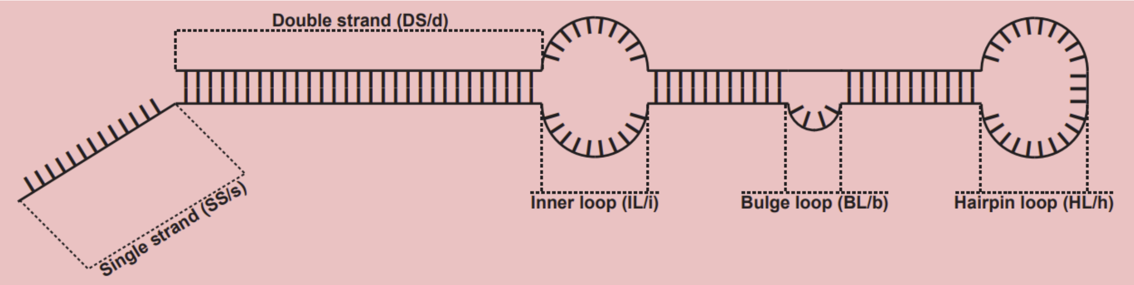


**Supplementary Methods Figure 2. Types of annotated secondary structure features.**

We also use the program full_premrna_not_edited_sequence in order to get the pre-mRNA sequences of those genes that don't have any edited adenosines. This program uses as input the same files as the full_premrna_sequence program and we use it with the same options, except for the -d option, because in the current version, this program doesn't accept it. It outputs a single file with the pre-mRNA sequences that don't contain any edited adenosine. We use the split command with the options -d and -n 61 to separate the sequence file in 61 files. For these non-edited sequences, we use linearfold, regroup the structure files with the cat command and the structure_features program in the same way as with those sequences that have editing.

We now use different programs to get the different datasets for the random forest and neural networks. For the random forest datasets, we need both positive and negative datasets (edited and non-edited adenosines). We use the program non_editing_adenosines to get the positions within the sequences of non-edited adenosines. This program uses as input the editing positions annotation file from the full_premrna_sequence (or an empty file for the sequences that have no editing) and the linearfold output with the secondary structure predictions (even if we only use the sequence) and it outputs the positions of all the adenosines in the pre-mRNA sequences that are not annotated as edited divided across 40 separate files.

We then use the program editing_structures_DB to get the files that will be used as datasets for the random forest algorithm (Supp. Met. Fig. 1). This program uses as input the edited (or non edited) positions files and the secondary structure features annotation file and it outputs a table where for each adenosine we have a series of descriptors about the secondary structure, both in the whole transcript, within a local window around the target adenosine (defined by the -s option), as well as details for a number (defined by the -m option) of the closest features of each type to the target adenosine. We always use -m 5, but for each file we run the program 4 times with the -s option for windows of 50, 200, 500 and 1000 nucleotides on both sides of the targeted adenosines. We do this for the single file with the edited positions and for all 80 files with non edited positions (40 for the sequences that have at least one editing and 40 for the sequences that have no editing). We then use the program random_lines to create the definitive negative datasets of the same size as the positive ones by taking random positions from all the

80 negative datasets. The random_lines program is used with the -h option which keeps the first line of the input files as a header and the -n option to define the number of lines we want in the output dataset. We also use random_lines to create smaller positive and negative datasets of 2000, 10000 and 100000 positive and negative cases.

For the neural networks approach, we use the program deep_learning_encoder to get the datasets. (Supp. Met. Fig. 1) This program uses as input the editing positions annotation file from the full_premrna_sequence (or an empty file for the sequences that have no editing), the linearfold output with the secondary structure predictions and the structure_features output file. The output is a dataset (Supp. Met. Fig. 3) with the same sequences and structures as in the linearfold structures file, but with two added channels for each sequence, one codifying the feature type that is annotated in that position, and the other codifying weather a position corresponds to an edited adenosine (2), a non-edited adenosine (1) or another nucleotide (0). Note that for the neural networks approach we don't use the linearfold secondary structure channel, even if it is present in the dataset file, but instead use our feature channel; this is because in the later pre-processing of the datasets, when we cut the sequences into windows, the dots and parenthesis format of presenting the secondary structures may present an information loss.

```
LOC105379585    LOC105379585
GTCGGAGCAGGGGCCGCGGGCGGCAGCTGACGGGCGGGCCCTTCCGGAG
...((((((.((((.((((((((((((((....)))((((.((..((..(
sssddddddiddddiddddddddddddddhhhhdddddddiddiiddiid
0000001001000000000000000010000100000000000000000010

LYPD5   LYPD5
accgaatatgtgttgtctttaaatacagagtatcgagacctcagaccctt
..................................((((.....((((( 
ssssssssssssssssssssssssssssssssddddiiiiiddddd
100011010000000000001110101010010001010000101000000
```

**Supplementary Methods Figure 3. Neural network input.** An example of the neural networks dataset before the last part of the processing.

Description of all the descriptors used in the random forest datasets

The descriptors used for the random forest datasets have been defined to try to capture most of the secondary structure information we thought could be relevant. In general terms, we have three types of descriptors (Supp. Met. Fig. 4):

- Global descriptors: these are statistics about each type of feature in the scope of the whole pre-mRNA sequence.
- Local descriptors: these are more or less the same statistics as in the global descriptors, within the scope of a local window defined by a number of nucleotides on both sides of

the edited adenosine. Note that if a feature lies partially inside the window it is still counted.

- XClosest descriptors: here we have more detailed information about concrete features in relation to the target adenosine. These descriptors are calculated for each type of structure, for a number (M = 5) of features closest to the target adenosine.

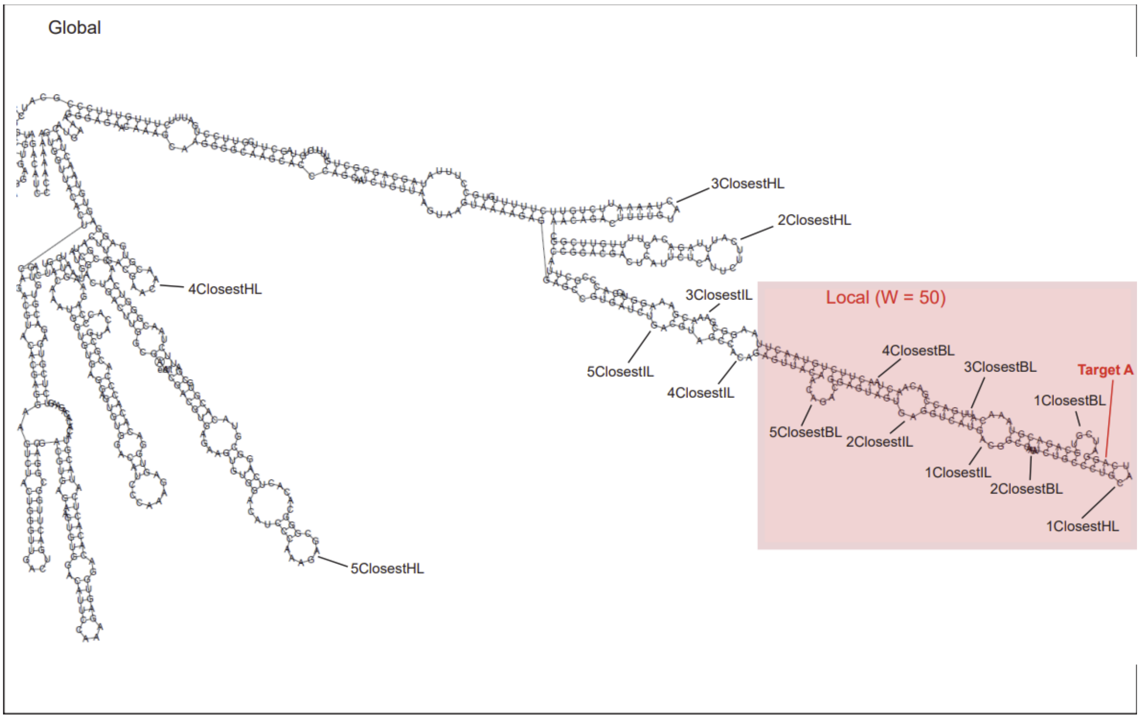


**Supplementary Methods Figure 4. Schematic representation of an RNA molecule with some of the structures used as descriptor**s. The local window (red square) provides the data for the local descriptors, while the global descriptors use the whole molecule. Target adenosine tagged in red. the Xclosest descriptors refer to the Xth feature of that type closest to the target adenosine, independently of the local window.

The following table (Supp. Met. Table 1) shows a description of all the descriptors (not that XClosest descriptors are variable in number, multiplied by M):

| Descriptor | Type | Description |
|---|---|---|
| GlobalPercDS | Global | The percentage of nucleotides that are in double strand (DS) in the whole pre-mRNA sequence |
| GlobalMaxDSSize | Global | The length in nucleotides of the largest DS fragment in the whole pre-mRNA sequence |
| GlobalNumHL | Global | A count of hairpin loops (HL) in the whole pre-mRNA sequence |

| | | |
|---|---|---|
| GlobalMaxHLSize | Global | The size in nucleotides of the largest HL in the whole pre-mRNA sequence |
| GlobalAvgHLSize | Global | The average size in nucleotides of all the HL in the whole pre-mRNA sequence |
| GlobalNumIL | Global | A count of inner loops (IL) in the whole pre-mRNA sequence |
| GlobalMaxILSize | Global | The size in nucleotides of the largest IL (counting in the strand where the size of the particular IL is larger) in the whole pre-mRNA sequence |
| GlobalAvgILSize | Global | The average size in nucleotides of all the IL (counting in the strand where the size of the particular IL is larger) in the whole pre-mRNA sequence in the whole pre-mRNA sequence |
| GlobalNumBL | Global | A count of bulge loops (BL) in the whole pre-mRNA sequence |
| GlobalMaxBLSize | Global | The size in nucleotides of the largest BL in the whole pre-mRNA sequence |
| GlobalAvgBLSize | Global | The average size in nucleotides of all the BL in the whole pre-mRNA sequence |
| EventInDS | Local | A binary descriptor (0 or 1), that tells whether the target adenosine is inside of a DS fragment |
| EventInNick | Local | A binary descriptor (0 or 1), that tells whether the target adenosine is inside of a nick (defined as an inner loop or a bulge loop of size 1) |
| LocalDSperc | Local | The percentage of nucleotides that are in DS in the local window |
| LocalDistToDS | Local | The distance in nucleotides from the target adenosine to its closest DS fragment (can be 0 if the target adenosine is inside a DS fragment) |

| | | |
|---|---|---|
| LocalClosestDSSize | Local | The length in nucleotides of the DS fragment closest to the target adenosine |
| LocalNumHL | Local | A count of HL in the local window |
| LocalMaxHLSize | Local | The size in nucleotides of the largest HL in the local window |
| LocalAvgHLSize | Local | The average size in nucleotides of all the HL in the local window |
| LocalNumIL | Local | A count of IL in the local window |
| LocalMaxILSize | Local | The size in nucleotides of the largest IL (counting in the strand where the size of the particular IL is larger) in the local window |
| LocalAvgILSize | Local | The average size in nucleotides of all the IL (counting in the strand where the size of the particular IL is larger) in the whole pre-mRNA sequence in local window |
| LocalNumBL | Local | A count of BL in the local window |
| LocalMaxBLSize | Local | The size in nucleotides of the largest BL in the local window |
| LocalAvgBLSize | Local | The average size in nucleotides of all the BL in the local window |
| LocalNumSS | Local | A count of a single strand (SS) fragment other than HL, BL or IL in the local window |
| LocalMaxSSSize | Local | The size in nucleotides of the largest SS fragment other than HL, BL or IL in the local window |
| LocalAvgSSSize | Local | The average size in nucleotides of all the BL in the local window |
| XClosestHLEventInside | XClosest | A binary descriptor (0 or 1) that tells whether the target adenosine is inside of the Xth closest HL to it (can only be true for the 1st closest) |

| | | |
|---|---|---|
| XClosestHLDistanceToEvent | XClosest | The distance in nucleotides from the target adenosine to the closest end of the Xth closest HL to it (this is the absolute value of XClosestHLClosestEndRelPos) |
| XClosestHLClosestEndRelPos | XClosest | The relative position in nucleotides (negative = 5’ direction, positive = 3’ direction) of the closest end of the Xth closest HL to the target adenosine in relation to it |
| XClosestHLMidRelPos | XClosest | The relative position in nucleotides (negative = 5’ direction, positive = 3’ direction) of the middle point of the Xth closest HL to the target adenosine in relation to it |
| XClosestHLSize | XClosest | The size in nucleotides of the Xth closest HL to the target adenosine |
| XClosestILEventInside | XClosest | A binary descriptor (0 or 1) that tells whether the target adenosine is inside of the Xth closest IL to it (can only be true for the 1st closest) |
| XClosestILIsNick | XClosest | A binary descriptor (0 or 1) that tells whether the Xth closest IL to the target adenosine is a nick (an IL of 1 nucleotide in size on both strands) |
| XClosestILDistanceToEvent | XClosest | The distance in nucleotides from the target adenosine to the closest end of the Xth closest HL to it (this is the absolute value of XClosestILClosestEventStrandEndRelPos) |
| XClosestILClosestEventStrandEndRelPos | XClosest | The relative position in nucleotides (negative = 5’ direction, positive = 3’ direction), within the same strand as the target adenosine, of the closest end of the Xth closest HL to the target adenosine in relation to it |
| XClosestILEventStrandMidRelPos | XClosest | The relative position in nucleotides (negative = 5’ direction, positive = 3’ direction), within the same strand as |

| | | |
|---|---|---|
| | | the target adenosine, of the middle point of the Xth closest HL to the target adenosine in relation to it |
| XClosestILEventStrandSize | XClosest | The size in nucleotides, corresponding to the same strand as the target adenosine, of the Xth closest HL to the target adenosine |
| XClosestILOppositeStrandMidRelPos | XClosest | The relative position in nucleotides (negative = 5' direction, positive = 3' direction), within the strand opposite to the target adenosine, of the middle point of the Xth closest HL to the target adenosine in relation to it |
| XClosestILOppositeStrandSize | XClosest | The size in nucleotides, corresponding to the strand opposite to the target adenosine, of the Xth closest HL to the target adenosine |
| XClosestBLEventInside | XClosest | A binary descriptor (0 or 1) that tells whether the target adenosine is inside of the Xth closest BL to it (can only be true for the 1st closest) |
| XClosestBLIsNick | XClosest | A binary descriptor (0 or 1) that tells whether the Xth closest BL to the target adenosine is a nick (an BL of 1 nucleotide in size) |
| XClosestBLDistanceToEvent | XClosest | The distance in nucleotides from the target adenosine to the closest end of the Xth closest BL to it (this is the absolute value of XClosestBLClosestEventStrandEndRelPos) |
| XClosestBLClosestEventStrandEndRelPos | XClosest | The relative position in nucleotides (negative = 5' direction, positive = 3' direction), within the same strand as the target adenosine, of the closest end of the Xth closest BL to the target adenosine in relation to it |

| | | |
|---|---|---|
| XClosestBLEventStrandMidRelPos | XClosest | The relative position in nucleotides (negative = 5’ direction, positive = 3’ direction), within the same strand as the target adenosine, of the middle point of the Xth closest BL to the target adenosine in relation to it |
| XClosestBLEventStrandSize | XClosest | The size in nucleotides, corresponding to the same strand as the target adenosine, of the Xth closest BL to the target adenosine (one of the two sizes of a BL will always be 0) |
| XClosestBLOppositeStrandMidRelPos | XClosest | The relative position in nucleotides (negative = 5’ direction, positive = 3’ direction), within the strand opposite to the target adenosine, of the middle point of the Xth closest BL to the target adenosine in relation to it |
| XClosestBLOppositeStrandSize | XClosest | The size in nucleotides, corresponding to the strand opposite to the target adenosine, of the Xth closest BL to the target adenosine (one of the two sizes of a BL will always be 0) |

**Supplementary Methods Table 1.** Reference table for all the descriptors used on the Random Forest datasets. Note that the descriptors of the XClosest type appear a number of times as defined by the M variable.

#### Starting Neural Networks dataset structure

The initial data are genomic sequences annotated with the secondary structures of the pre-mRNA and with the editing positions. The initial data file is generated, which contains all the pre-mRNA sequences along with the prediction of their secondary structure and the annotations of the edited adenosine positions. The file is encoded for each gene as follows (Supp. Met. Fig. 5):

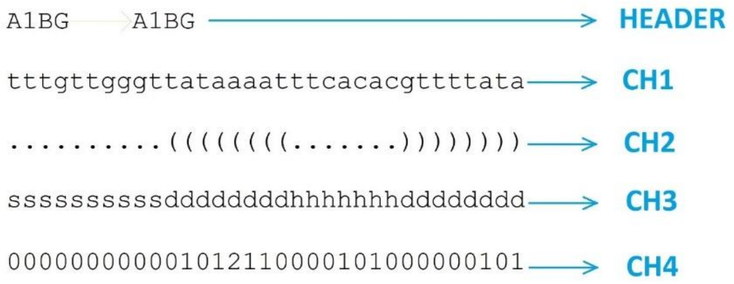


**Supplementary Methods Figure 5. Original encryption of the data.**

- HEADER: is the FASTA header of the gene sequence.
- CH1 (DNA): is the sequence of DNA AGCT nucleotides that encode the pre-mRNA of the gene.
- CH2 (STRUCTURE-PAIRS): is the representation of paired nucleotides represented by parentheses. The points would be unpaired nucleotides and the open and closed parentheses would be paired, following the order of pairing of the left parenthesis closest to the right parenthesis on its left side. It is known as "dot-bracket" notation because it uses the symbols ".", "(" and ")".
- CH3 (STRUCTURE-FEATURES): represents the belonging of each nucleotide to a substructure or special geometric motif in the pre-mRNA, which we call features of the structure. These motifs have been annotated with letters according to the following encoding:
  - **s** = single strand
  - **d** = double strand
  - **h** = hairpin loop
  - **i** = indown loop
  - **b** = bulge loop
- CH4: represents whether the adenosine nucleoside, in that position, is editable (value=2) or not (value=1). If there is no adenosine in that position it is represented by a zero (value=0).

### Window sequences

The genomic data files had to be processed properly in order to train the deep learning models. For this purpose, we have used fixed length window sequences, 2n+1, centered on each of the target adenosines for a given gene. The length of these sequences was also a hyperparameter that had to be adjusted. Window sequences of different lengths were tested, finding that the best results were obtained for n = 50, generating windows of 50+1+50 nucleotides. This size is long enough to find a clear learning signal, but short enough for computing times be reasonably short.

This cut must be made identically in all channels (CH1 to CH3). The code indicating whether there is an edit (value=2) or not (value=1) is extracted from the CH4 channel. The values “0” correspond to positions in which there is no Adenosine. The CH4 channel provides therefore the class category. (Supp. Met. Fig. 6)

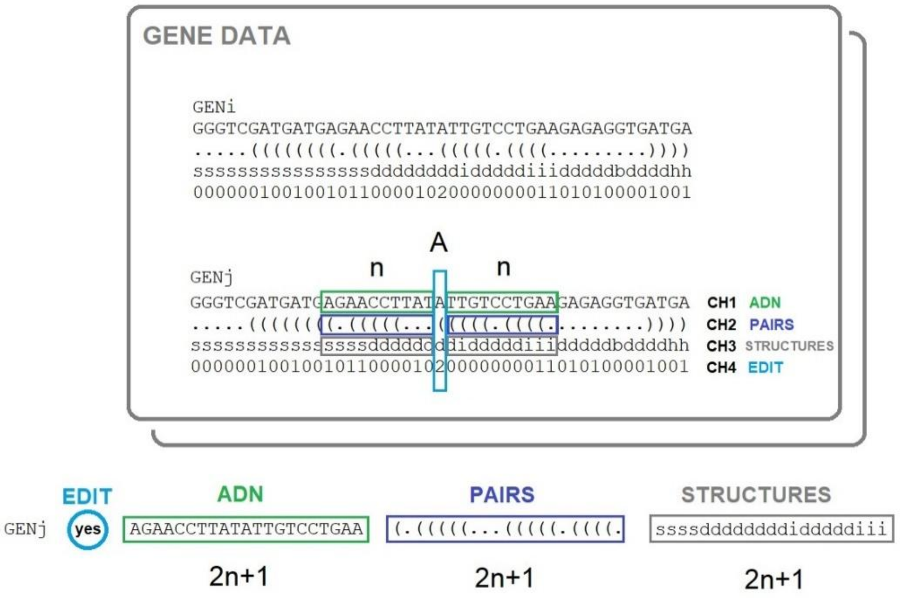


**Supplementary Methods Figure 6. Basic scheme of cutting the sequences**.

### Data encoding

Each nucleotide position originally had three information channels: CH1 (DNA), CH2 (PAIRS) and CH3 (FEATURES). We have tested different coding models and, in the end, the most efficient model has turned out to be the one that encodes only with the channels of primary sequence of nucleotides (CH1) and secondary substructures (CH3), which has allowed us to perform a simple coding of dimension 21, corresponding to 4x5 =20 symbols, for the two channels plus 1 additional symbol (*) for the *padding* of adenosine windows that are in extreme positions and cannot form symmetrical windows. While, originally, the CH3 is directly derived from CH2, when cutting into windows, the dot and parenthesis annotation of the CH2 has an information loss. Because one position within the window can be paired with a position outside the window, which would make the deduction of the corresponding features impossible once cut into windows. In order to preserve this information, we used CH3 instead of CH2.

Finally, the resulting 21 symbols were coded using one-hot notation (Supp. Met. Table 2).

| Tuple encoding | One-Hot Coding. |
|---|---|
| ('A', 's') / 0 | 0 / 100000000000000000000 |
| ('A', 'd') / 1 | 1 / 010000000000000000000 |
| ('A', 'h') / 2 | 2 / 001000000000000000000 |
| ('A', 'i') / 3 | 3 / 000100000000000000000 |
| ('A', 'b') / 4 | 4 / 000010000000000000000 |
| ('G', 's') / 5 | 5 / 000001000000000000000 |

| | |
|---|---|
| ('G', 'd') / 6 | 6 / 000000100000000000000 |
| ('G', 'h') / 7 | 7 / 000000010000000000000 |
| ('G', 'i') / 8 | 8 / 000000001000000000000 |
| ('G', 'b') / 9 | 9 / 000000000100000000000 |
| ('C', 's') / 10 | 10 / 000000000010000000000 |
| ('C', 'd') / 11 | 11 / 000000000001000000000 |
| ('C', 'h') / 12 | 12 / 000000000000100000000 |
| ('C', 'i') / 13 | 13 / 000000000000010000000 |
| ('C', 'b') / 14 | 14 / 000000000000001000000 |
| ('T', 's') / 15 | 15 / 000000000000000100000 |
| ('T', 'd') / 16 | 16 / 000000000000000010000 |
| ('T', 'h') / 17 | 17 / 000000000000000001000 |
| ('T', 'i') / 18 | 18 / 000000000000000000100 |
| ('T', 'b') / 19 | 19 / 000000000000000000010 |
| ('*', '*') / 20 | 20 / 000000000000000000001 |

**Supplementary Methods Table 2.** One-hot encoding of streams.

On the left side of the table there is the encoding of the tuples in the form of integers. And on the right side is the one-hot encoding. The integer encoding is more compact and readable than one-hot and is the one we save in a csv file. One-hot coding is done "on the fly", at runtime, during the training and validation of the neural network and during the testing of the sequences.

**Recurrent neural networks and global attention layer**

In this work we have used recurrent neural networks (RNN), first of all because this type of network is specially oriented to the processing of sequential data, such as genomic sequences.

Secondly, we also know that the nucleotides of the sequences can present relationships between distant positions in the primary sequence, since they can be geometrically close, because of the folding of the secondary structures. This feature also advises the use of recurrent networks, since they can better capture those relationships between distant positions in the sequence than a convolutional network, for example.

In particular we have implemented an LSTM (Long short-term memory) (Hochreiter & Schmidhuber, 1997). We have used a bidirectional LSTM, which is composed of two parallel layers that process the sequences in opposite directions. Bidirectional networks tend to offer better performance than unidirectional LSTMs and treat data in a symmetrical way, with respect to the two possible directions of reading. We have also added a layer of global attention (Supp. Met. Fig. 7).

The attention layer is a mechanism that allows the output of the recurrent network don't depend only on last hidden state but learn to give different weights to different hidden states to generate the output, seeking to give more relevance to the positions that are more decisive when classifying the sequence.

To develop the recurring deep learning models we have used the implementation of LSTM that makes tensorflow.keras, through the classes layers.Bidirectional and layers.LSTM (Supp. Met. Fig. 7 and Supp. Met. Fig. 8).

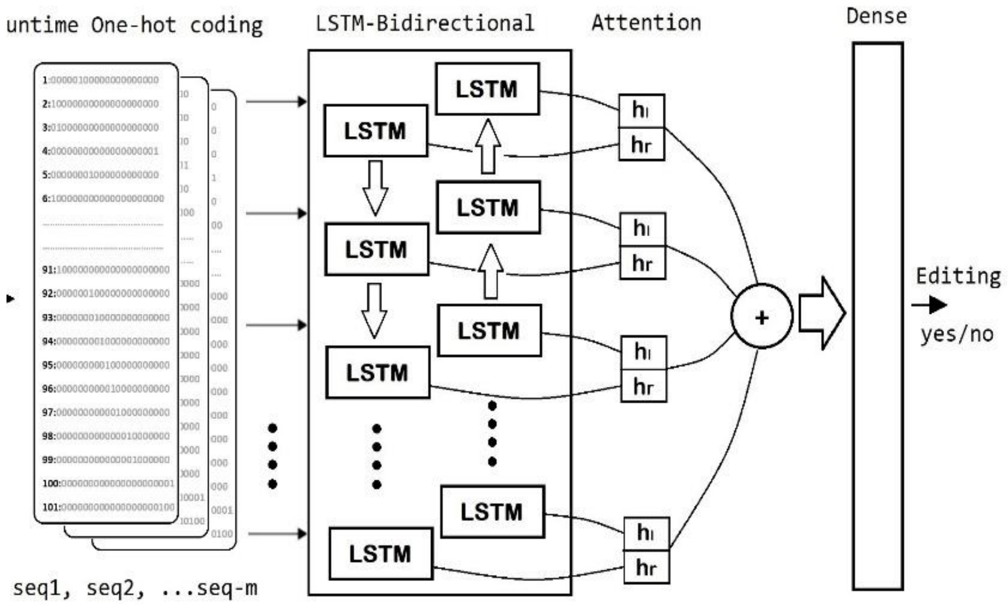


**Supplementary Methods Figure 7. Basic scheme of biLSTM.**

Model: "TFM_LSTMBidiAttention"

| Layer (type) | Output Shape | Param # | Connected to |
|---|---|---|---|
| INPUTS (InputLayer) | [(None, 101, 21)] | 0 | [] |
| LSTM_BI (Bidirectional) | (None, 101, 512) | 569344 | ['INPUTS[0][0]'] |
| dropout (Dropout) | (None, 101, 512) | 0 | ['LSTM_BI[0][0]'] |
| dense (Dense) | (None, 101, 1) | 513 | ['dropout[0][0]'] |
| flatten (Flatten) | (None, 101) | 0 | ['dense[0][0]'] |
| activation (Activation) | (None, 101) | 0 | ['flatten[0][0]'] |
| repeat_vector (RepeatVector) | (None, 512, 101) | 0 | ['activation[0][0]'] |
| permute (Permute) | (None, 101, 512) | 0 | ['repeat_vector[0][0]'] |
| multiply (Multiply) | (None, 101, 512) | 0 | ['LSTM_BI[0][0]', 'permute[0][0]'] |
| lambda (Lambda) | (None, 512) | 0 | ['multiply[0][0]'] |
| OUTPUT (Dense) | (None, 1) | 513 | ['lambda[0][0]'] |

Total params: 570,370
Trainable params: 570,370
Non-trainable params: 0

**Supplementary Methods Figure 8. Summary of the implemented biLSTM network.**

### Balancing training data

In its native state, the number of editable adenosines is very small regarding to the total number of adenosines. For example, in humans, the number of edited adenosines barely exceeds 2.7% of the whole number of adenosines. This fact makes our native training sets strongly unbalanced, presenting many more negative cases (non-editable) than positive (editable). As sufficient positive samples are available, it has been decided to address the problem of imbalance by means of sample balancing, in which all positive samples are taken and balanced by the same number of negative ones. Therefore, since the use of class weights failed to compensate for the disproportion between classes, we have balanced positive and negative samples in a 1 to 1 ratio.

### Dataset sizes

In the following table (Supp. Met. Table 3-5) we list the initial and final dataset sizes for all the training and testing datasets.

| **Species** | **Editing DB size** | **Editing NN training** | **Non-editing NN training** | **Editing NN testing** | **Non-editing NN testing** |
|---|---|---|---|---|---|
| *H. sapiens* | 15681871 | 107393 | 107410 | 22969 | 23061 |
| *H. sapiens* (old) | 4668508 | 40368 | 40456 | 8650 | 8670 |
| *M. musculus* | 107094 | 20880 | 20625 | 4386 | 4509 |
| *T. trachurus* | N/A | 99118 | 98990 | 21117 | 21335 |
| *O. bimaculoides* | N/A | 88781 | 88692 | 7855 | 7805 |

**Supplementary Methods Table 3.** Initial database sizes and final Neural Networks dataset sizes. Note that the initial database sizes are before any processing, as they appear in the REDIportal databases. For *Trachurus trachurus* and *Octopus bimaculoides* we didn't use the REDIportal databases.

| **Species** | **Editing RF W=50 training** | **Non-editing RF W=50 training** | **Editing RF W=50 testing** | **Non-editing RF W=50 testing** | **Editing RF W=200 training** | **Non-editing RF W=200 training** | **Editing RF W=200 testing** | **Non-editing RF W=200 testing** |
|---|---|---|---|---|---|---|---|---|

| | | | | | | | | |
|---|---|---|---|---|---|---|---|---|
| *H. sapiens* | 6648 | 6592 | 3336 | 3284 | 6259 | 5685 | 3097 | 2875 |
| *M. musculus* | 6413 | 5619 | 3224 | 2793 | 6316 | 6073 | 3174 | 3021 |

**Supplementary Methods Table 4.** Final dataset sizes for Random Forest with windows of 50 and 200.

| Species | Editing RF W=500 training | Non-editing RF W=500 training | Editing RF W=500 testing | Non-editing RF W=500 testing | Editing RF W=1000 training | Non-editing RF W=1000 training | Editing RF W=1000 testing | Non-editing RF W=1000 testing |
|---|---|---|---|---|---|---|---|---|
| *H. sapiens* | 6608 | 6510 | 3340 | 3220 | 6644 | 6587 | 3339 | 3277 |
| *M. musculus* | 6620 | 6595 | 3325 | 3283 | 6627 | 6663 | 3344 | 3302 |

**Supplementary Methods Table 5.** Final dataset sizes for Random Forest with windows of 500 and 1000.
